# Structural basis of LhcbM5-mediated state transitions in green algae

**DOI:** 10.1101/2021.03.02.433643

**Authors:** Xiaowei Pan, Ryutaro Tokutsu, Anjie Li, Kenji Takizawa, Chihong Song, Kazuyoshi Murata, Tomohito Yamasaki, Zhenfeng Liu, Jun Minagawa, Mei Li

## Abstract

In green algae and plants, state transitions serve as a short-term light acclimation process to regulate light harvesting capacity of photosystems I and II (PSI and PSII). During the process, a portion of the light-harvesting complexes II (LHCII) are phosphorylated, dissociate from PSII and bind PSI to form PSI-LHCI-LHCII supercomplex. Here we report high-resolution structures of PSI-LHCI-LHCII supercomplex from *Chlamydomonas reinhardtii*, revealing the mechanism of assembly between PSI-LHCI complex and two phosphorylated LHCII trimers containing all four types of LhcbM proteins. Two specific LhcbM isoforms, namely LhcbM1 and LhcbM5, directly interact with the PSI core through their phosphorylated amino-terminal regions. Furthermore, biochemical and functional studies on mutant strains lacking either LhcbM1 or LhcbM5 indicate that only LhcbM5 is indispensable in the supercomplex formation. The results unraveled the specific interactions and potential excitation energy transfer routes between green algal PSI and two phosphorylated LHCIIs.

## Introduction

During oxygenic photosynthesis, light-harvesting complexes (LHCs) absorb photon energy and supply excitation energy for subsequent charge separation processes occurring in photosystems I and II (PSI and PSII). In the green lineage (green algae and plants), both photosystems consist of two major modules: the core complex containing the cofactors for charge separation, electron transport and light harvesting processes, and the outer LHCs extending the light absorption capacity of the core ^1,2^. Most of the LHC proteins adopt similar folding with three transmembrane helices (TMHs) named helices B, C and A ^3^. LHCIs with apoproteins encoded by the *Lhca* genes are associated with the PSI core, forming the PSI-LHCI complex. LHCIIs with the apoproteins encoded by *Lhcb* genes are mostly attached to the PSII core, constituting the PSII-LHCII complex ^1^. Green algae contain more *Lhca* and *Lhcb* genes and larger antenna size for both photosystems in comparison with plants ^1,4–6^. In the unicellular green alga *Chlamydomonas reinhardtii*, there are nine *Lhca* genes (*Lhca* 1–9) ^5^ and up to ten LHCI monomers associated with the PSI core ^7,8^. For PSII antennae, 11 *Lhcb* genes encode apoproteins of nine LhcbM isoforms (LhcbM1–9, presumably forming the major trimeric LHCII) and two monomeric Lhcb proteins known as CP26 (Lhcb5) and CP29 (Lhcb4) ^9^. The nine LhcbM isoforms can be classified into four types, namely type I (LhcbM3, LhcbM4, LhcbM6, LhcbM8 and LhcbM9), type II (LhcbM5), type III (LhcbM2 and LhcbM7) and type IV (LhcbM1) ^9,10^. The five isoforms from type I show high sequence homology with each other, and the two type III isoforms share identical sequence for their mature polypeptides (hereafter referred to as LhcbM2/7).

PSI and PSII have different pigment compositions, and hence exhibit distinct absorption characteristics. Specifically, PSI has a broad absorption peak in the far-red region, whereas PSII favors red light ^11^. An imbalance of energy distribution between the two photosystems tends to occur in natural environments, where light quality and quantity fluctuate constantly ^12^. To ensure optimal photosynthetic efficiency, plants and algae have developed both short-term and long-term adaptation in response to the fluctuating illumination. State transitions serve as a short-term light acclimation mechanism to balance energy distribution between two photosystems ^11,12^. The process includes two distinct states (states 1 and 2) and involves reversible phosphorylation of LHCII and migration of mobile LHCIIs between PSI and PSII. The transition from state 1 to state 2 is facilitated by phosphorylation of the amino-terminal region of LHCII and association of phosphorylated LHCII with PSI to form the PSI-LHCI-LHCII supercomplex (also known as state 2 supercomplex). In the reverse process, LHCII is dephosphorylated by the state transition phosphatases and dissociates from PSI ^11–13^. In green algae, state transitions show stronger amplitude and play a wider photoprotective role than the process in plants^14^. It was demonstrated that only a limited fraction (20-25%) of LHCIIs in higher plants are involved in state transitions, whereas 35-38% of LHCIIs are involved in *C. reinhardtii*^15^. State transitions were suggested to alter the thylakoid ultrastructure and protein-protein interactions in *C. reinhardtii* ^16^. The process may also lead to switch of the detached LHCIIs into the energetically quenched form so as to avoid photodamage ^16–18^.

Previously, the high resolution structure of plant state 2 supercomplex revealed that one LHCII trimer stably binds to PSI core ^19^. Within the trimer, Lhcb2 is phosphorylated at its third residue threonine, and the first three residues directly interact with PSI core subunits PsaH, PsaL and PsaO. The state 2 supercomplex in *C. reinhardtii* is larger and more intricate compared to the one from plants. Previous biochemical analysis and a two-dimensional projection map of the state 2 supercomplex from *C. reinhardtii* suggested that two LHCII trimers and one monomer (likely CP29) are associated with the PSI core on the opposite side of the “LHCI belt” ^20–22^. Earlier reports indicated that LhcbM5 and LhcbM2/7 participate in the formation of state 2 supercomplex, whereas the absence of LhcbM1 does not affect state transitions but causes a decrease of thermal dissipation instead ^4,23^. Besides, the other work showed that PSI-LHCI-LHCII supercomplex in *C. reinhardtii* contains all four types of LhcbM proteins ^22^. Except LhcbM2/7, all other LhcbM proteins are phosphorylated under state 2 ^22,24^. Recently, it was found that two phosphatases, namely Protein Phosphatase1 (PPH1) and Photosystem II Core Phosphatase (PBCP), are both required for the transition from state 2 to state 1 in *C. reinhardtii* ^13^. The former is essential for dephosphorylation of LhcbM1 and LhcbM5, while the latter may dephosphorylate LhcbM5 and type I isoforms.

So far, the detailed structure of state 2 supercomplex in *C. reinhardtii* and the subunit composition of LHCII trimers associated with PSI are still unknown. Fundamental questions, as to which LhcbM isoforms bind to green algal PSI under state 2 and how two LHCII trimers establish specific interactions with PSI, remains unresolved. The potential excitation energy transfer (EET) pathways between LHCII and PSI-LHCI are to be unraveled at high resolution. Here we report single-particle cryo-electron microscopy (cryo-EM) structures of PSI-LHCI-LHCII supercomplexes from *C. reinhardtii* (*Cr*PSI-LHCI-LHCII). Our work provides structural and functional insights into the assembly mechanism of *Cr*PSI-LHCI-LHCII supercomplexes. The interfacial pigment-pigment connections between LHCIIs and PSI-LHCI complex, essential for understanding the molecular mechanism of energy distribution to PSI during state transitions in *C. reinhardtii*, have been revealed at near atomic resolution.

## Results

### Overall structure of *Cr*PSI-LHCI-LHCII

We purified *Cr*PSI-LHCI-LHCII supercomplexes from both wild type (WT) and the double phosphatase mutant *pph1;pbcp* ^13^ in which both PPH1 and PBCP were mutated and the cells are locked in state 2 (Supplementary Fig. 1), and solved their cryo-EM structures at 4.23 and 2.84 Å resolution, respectively (Supplementary Figs. 2, 3). The supercomplex prepared from the *pph1;pbcp* mutant is more stable and contains more intact target supercomplex particles. In addition, it represents the native *Cr*PSI-LHCI-LHCII supercomplex from WT strain because their components are essentially the same and their structures are nearly identical. Thus the 2.84 Å structure was further refined and the final model contains 29 subunits (including 13 PSI core subunits, 10 LHCI monomers and 2 LHCII trimers), 334 chlorophylls (Chls), 87 carotenoids, 2 phylloquinone, 3 Fe_4_S_4_ cluster and numerous lipids (Supplementary Fig. 4, Supplementary Tables 1 and 2). Eight LHCI proteins make two antenna belts (inner, Lhca3-Lhca7-Lhca8-Lhca1; outer, Lhca5-Lhca6-Lhca4-Lhca1) binding to the PsaK-PsaJ-PsaF-PsaG side, identical to the previously reported *Cr*PSI-LHCI structures^25,26^. Meanwhile, the Lhca9, Lhca2 and two LHCII trimers together form an additional antenna belt at the opposite side, attaching to PsaG-PsaI-PsaH-PsaL-PsaO (Fig. 1a). One LHCII trimer (LHCII-1) in *Cr*PSI-LHCI-LHCII is located at a position similar to the LHCII trimer in plant PSI-LHCI-LHCII supercomplex, but shows a rotational shift (Fig. 1b,c). The other LHCII trimer (LHCII-2) is specific to *Cr*PSI-LHCI-LHCII, and bridges the gap between LHCII-1 trimer and Lhca2.

**Figure 1.**
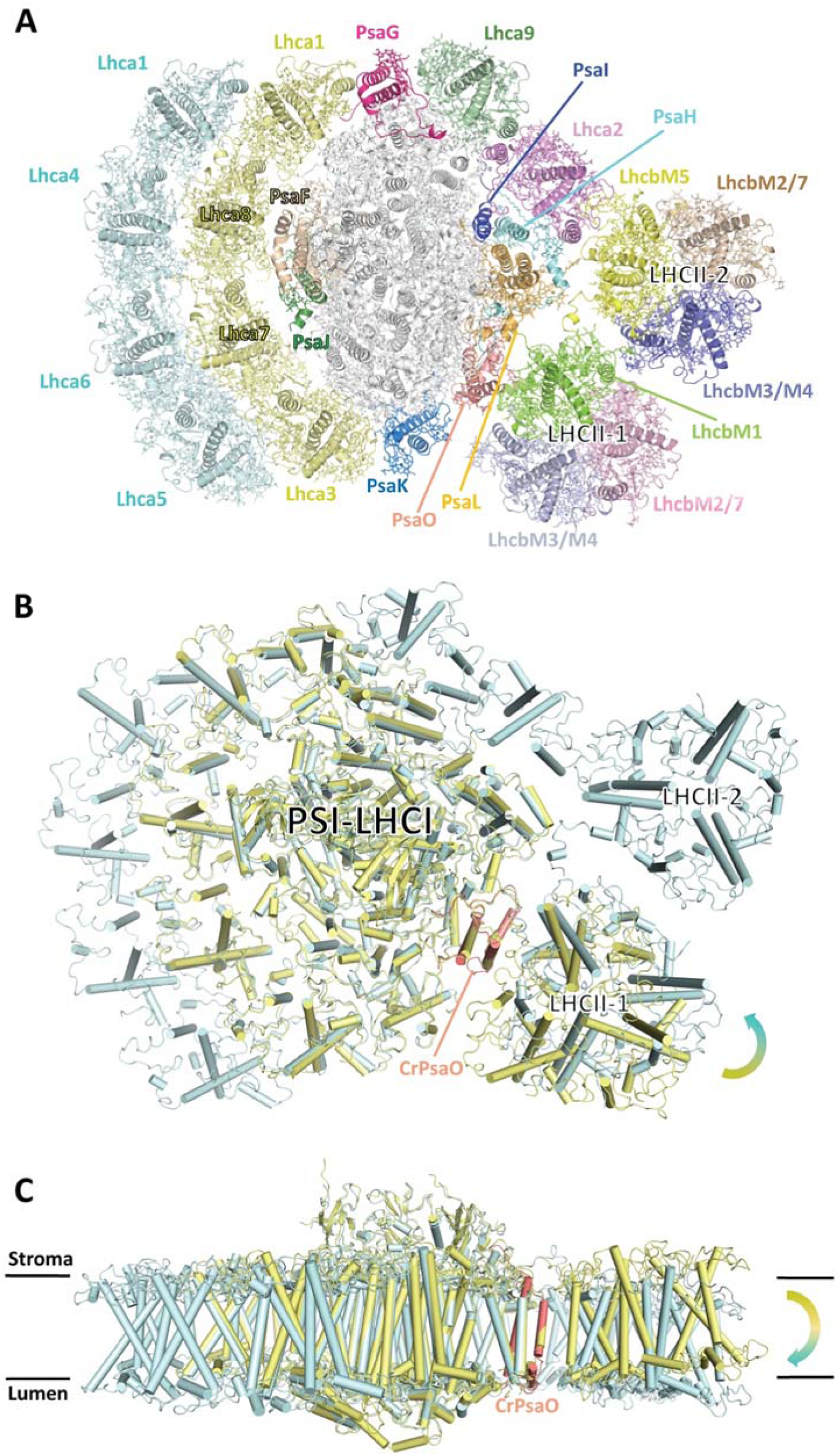
Overall structure of native *Cr*PSI-LHCI-LHCII supercomplex. (a) Cartoon representation of *Cr*PSI-LHCI-LHCII supercomplex viewed from the stromal side. Pigments, lipids and other cofactors are shown in stick mode. The core subunits PsaA-PsaE are shown in white, while other subunits are shown in different color and labeled. (b,c) Structural comparison of *Cr*PSI-LHCI-LHCII supercomplex (pale-cyan) with maize PSI-LHCI-LHCII supercomplex (yellow, PDB code 5ZJI), viewed from the stromal side (b) and from the membrane plane (c). *Cr*PsaO is highlighted in salmon and indicated. Arrows indicate the rotational direction of LHCII-1.

Basing on the high-quality cryo-EM density and the sequence characteristics of LhcbM proteins, we have identified the individual LhcbM proteins in each LHCII trimer (Supplementary Fig. 5). LHCII-1 is composed of LhcbM1, LhcbM2/7 and LhcbM3 (or LhcbM4, hereafter referred to as LhcbM3/M4, since the two isoforms in Type I cannot be further distinguished according to the present map), whereas LHCII-2 is composed of LhcbM5, LhcbM2/7 and LhcbM3/M4 (Fig. 1a). In our *Cr*PSI-LHCI-LHCII structure, all four types of LHCII associate with PSI under state 2 condition, consistent with the previous biochemical analysis ^22^. Although earlier reports suggested that monomeric CP29 may be associated with PSI in state 2, we did not observe any extra densities corresponding to it. It is possible that CP29 detected in the state 2 supercomplex reported previously may arise from the PSII contamination (Supplementary Figs. 1-3).

### Assembly between two LHCII trimers and PSI-LHCI complex

In the *Cr*PSI-LHCI-LHCII structure, two LhcbM proteins facing the PSI core were identified as LhcbM1 and LhcbM5 in LHCII-1 and LHCII-2, respectively (Fig. 1a, Supplementary Fig. 5). Both LhcbM proteins have a phosphorylated threonine (pThr) with well defined cryo-EM densities at their N-terminal tails (Supplementary Figs. 5a, 5d). The phosphorylation sites of LhcbM1 and LhcbM5 are consistent with the previous mass spectrometry analysis result ^24^. The phosphate groups of pThr residues have crucial role for the formation of state 2 supercomplex by forming salt bridges and hydrogen bonds with nearby residues (Fig. 2).

**Figure 2.**
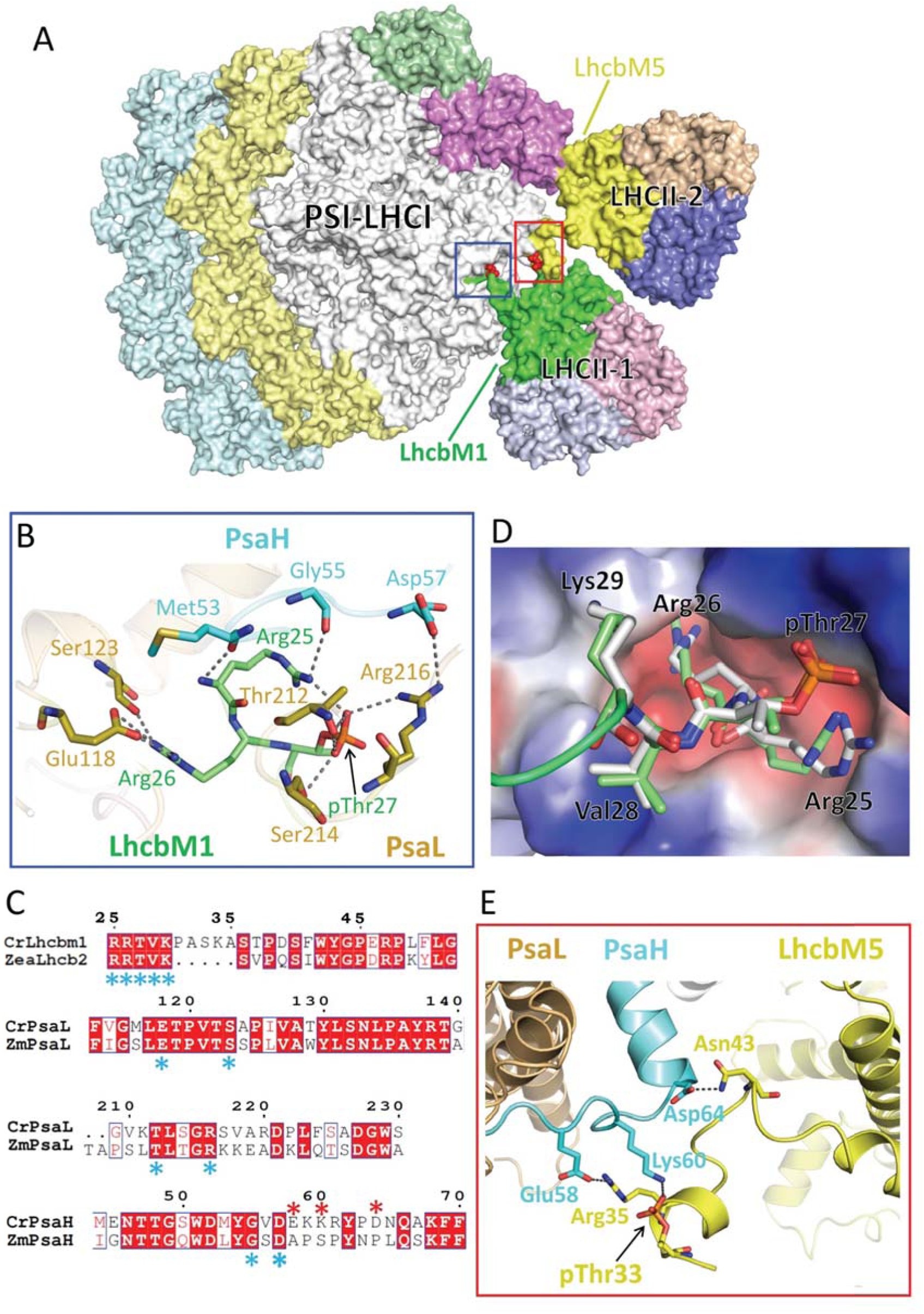
Assembly of two LHCII trimers with the PSI-LHCI. (a) Surface representation of CrPSI-LHCI-LHCII supercomplex with the pThr from LhcbM1 and LhcbM5 are highlighted in red. (b) The detailed interaction between the N-terminal region of LhcbM1 and the PSI core subunits PsaH and PsaL. (c) Sequence alignment of the interaction-related regions of LhcbM1, PsaL and PsaH from *C. reinhardtii* (Cr) with Lhcb2, PsaL and PsaH from *Zea mays* (Zm), respectively. The conserved and variant residues involved in the inter-subunit interactions are indicated by blue and red asterisk, respectively. The conserved and variant residues are indicated by blue and red asterisk, respectively. (d) Structural comparison of the N-terminal tail of *Cr*LhcbM1 and *Zm*Lhcb2. The PSI core is shown in electrostatic surface mode with red and blue representing acidic and basic regions, respectively. (e) The detailed interaction between the N-terminal region of LhcbM5 and the PSI core subunit PsaH. Hydrogen bond interactions are shown as black dashed lines.

In LHCII-1, the phosphorylated N-terminal region of LhcbM1 is rope-shaped and reaches out to the surficial pocket on PSI (Fig. 2a). Residue pThr27 and two preceding basic residues in LhcbM1 form specific interactions with the core subunits PsaH and PsaL (Fig. 2b). This recognition and interacting pattern resembles those observed in plant state 2 supercomplex, with LhcbM1 functioning similarly as Lhcb2 in plants. In line with the structural observation, LhcbM1 is the only LhcbM isoform in *C. reinhardtii* that possesses an N-terminal motif with amino acid sequence (^25^RRtVK^29^) identical to plant Lhcb2 (Fig. 2c). These five residues of LhcbM1 and plant Lhcb2, including the phosphate group, superpose well in the two structures (Fig. 2d). In addition, the residues from PsaH and PsaL interacting with LhcbM1 are invariant between plants and *C. reinhardtii* (Fig. 2c), suggesting that the recognition between the phosphorylated LHCII-1 trimer and the PSI core is conserved in the green lineage. However, when compared with the LHCII trimer in plant PSI-LHCI-LHCII, LHCII-1 in *Cr*PSI-LHCI-LHCII rotates around its N-terminal tail within the membrane plane towards LHCII-2 and pivots towards the lumen by approximately 10° and 15°, respectively (Fig. 1b,c). The rotation results in weaker association between LhcbM1 and the core subunit PsaO compared to plant Lhcb2-PsaO interactions.

The LHCII-2 trimer found in *Cr*PSI-LHCI-LHCII is absent in plant state 2 supercomplex and it associates with PSI and Lhca2 through LhcbM5 (Fig. 2a). Previously, LhcbM5 was found to be phosphorylated under state 2 conditions and present in the *Cr*PSI-LHCI-LHCII supercomplex ^21,27^. Compared with other LhcbM proteins, LhcbM5 has an extended N-terminal region (Supplementary Fig. 5d), forming a short α-helix and a loop intercalating into the gap between PsaH-PsaL and LhcbM1 of the LHCII-1 trimer (Figs. 2a, 2e). The phosphate group of pThr33 of LhcbM5 binds to Lys60 from PsaH through strong electrostatic interaction. Following pThr33, residues Arg35 and Asn43 of LhcbM5 form salt bridge and hydrogen bond with Glu58 and Asp64 from the stromal loop of PsaH (Fig. 2e). The three residues from PsaH involved in binding LhcbM5 are variable in maize PsaH (Fig. 2c), suggesting that the PSI docking site for LHCII-2 is specific to green algae and not conserved in plants.

While LhcbM5 binds PsaH through its N-terminal tail, it also interacts extensively with Lhca2 (Fig. 3a-c). The AC loop and BC loop (the loop connecting helices A and C, and helices B and C) of LhcbM5 and Lhca2 contact each other and form multiple interactions at stromal and luminal sides, respectively (Fig. 3b, c). Furthermore, one lipid molecule (MGDG) and several interfacial pigments at the luminal side stabilize the association of LhcbM5 with Lhca2 through hydrophobic interactions (Fig. 3c).

**Figure 3.**
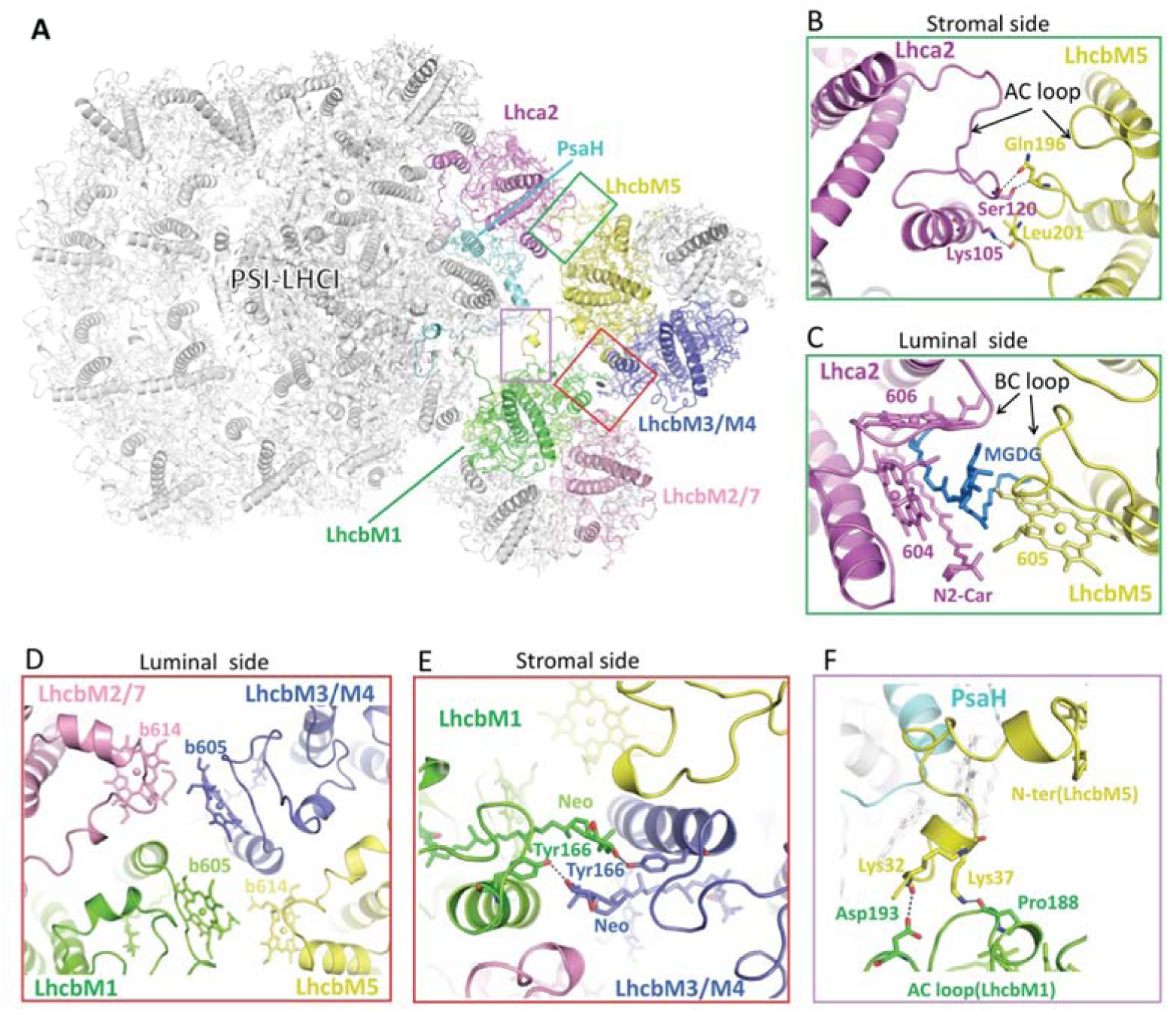
Interaction between two LHCII trimers and between LHCII and Lhca2. (a) Cartoon representation of *Cr*PSI-LHCI-LHCII supercomplex. LhcbM1 and LhcbM2/7 from LHCII-1, LhcbM5 and LhcbM3/M4 from LHCII-2, Lhca2 and PsaH are highlighted in colors same as in Fig. 1a, while other subunits are shown in white. Two LHCII trimers are related by a pseudo-C2 axis which is indicated by an elliptical ring. (b,c) The detailed interaction between LhcbM5 and Lhca2 at the stromal (b) and luminal (c) side, respectively. One lipid molecule monogalactosyldiacyl glycerol (MGDG, blue) serves as a “molecular glue” linking LhcbM5 and Lhca2 at the luminal side. (d) Hydrophobic interaction between two LHCII trimers at the luminal side. (e) Hydrogen bond interactions between neoxanthin (Neo) and Tyr166 of both LhcbM1 and LhcbM3/M4 from LHCII-2. (f) The detailed interaction between the N-terminal region of LhcbM5 and the AC loop of LhcbM1. Residues, pigments and lipids involved in the inter-subunit interactions are labeled and shown in stick mode, with the central-Mg of chlorophylls shown as spheres.

The two LHCII trimers are related with each other through a pseudo-C2 axis running through their interface (Fig. 3a). They interact closely with each other mainly through hydrophobic interactions between interfacial pigments from LhcbM1 and LhcbM2/7 of LHCII-1 and from LhcbM5 and LhcbM3/M4 of LHCII-2 (Fig. 3d). Moreover, LhcbM1 and LhcbM3/M4 of LHCII-2 form mutual hydrogen bond interactions between their neoxanthin molecule and Tyr166 at the stromal side (Fig. 3e). Furthermore, two lysine residues (Lys32 and Lys37) in the N-terminal extended region of LhcbM5 are hydrogen bonded to the stromal AC loop of LhcbM1 (Fig. 3f). The rotational shift of LHCII-1 trimer in *Cr*PSI-LHCI-LHCII supercomplex (relative to the one in plant PSI-LHCI-LHCII supercomplex) facilitates the mutual interactions between two LHCII trimers and thus stabilizes them within the additional antenna belt (Fig. 1b).

### LhcbM5, but not LhcbM1, is essential for state 2 supercomplex formation

Evidently, LHCII-2 is located at a central position in the state 2 supercomplex, as the phosphorylated LhcbM5 interacts with the PSI core (through PsaH), Lhca2 and LHCII-1 trimer simultaneously (Fig. 1a). In the *ΔLhcbM5* (lacking LhcbM5) mutant (Supplementary Fig. 6), the PSI-LHCI-LHCII supercomplex disappeared almost completely (Fig. 4a-c), confirming that LhcbM5 has a crucial role in the formation of state 2 supercomplex. To investigate the state transition phenotype of the *ΔLhcbM5* mutant, we further evaluated its qT quenching capability under the light conditions for induction of state transitions (Fig. 4d). Both WT and the *ΔLhcbM5* mutant strains showed qT quenching, a decrease of maximum chlorophyll fluorescence (Fm), under state 2-inducing blue light. Remarkably, the *ΔLhcbM5* mutant demonstrated significantly slower transition from state 1 to state 2 upon induction by blue light, whereas there was little difference in the state 2 to state 1 fluorescence kinetics between the two strains (Fig. 4d). Therefore, both structural finding and the *in vivo* analysis of qT quenching capability of the *ΔLhcbM5* mutant underscore the physiological importance of LhcbM5 during state transitions in green algae.

**Figure 4.**
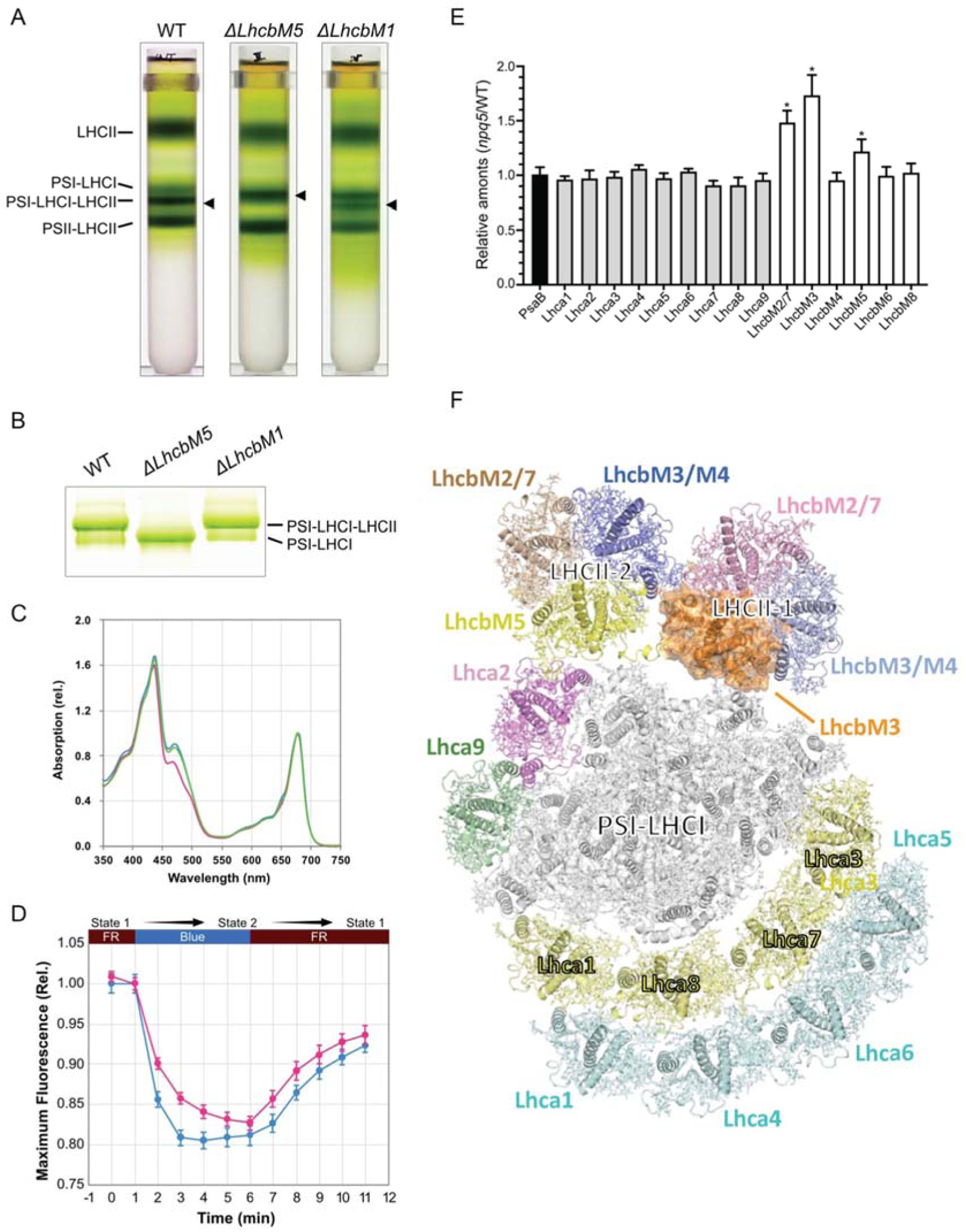
Purification and characterization of the supercomplex samples from *ΔLhcbM1* and *ΔLhcbM5* mutants. (a) Sucrose gradient fractionation of solubilized state 2 thylakoid membranes from WT, *ΔLhcbM1*, and *ΔLhcbM5* mutants. Each band shown is labeled. (b) CN-PAGE analysis of purified PSI-LHC(-LHCII) supercomplexes from WT, *ΔLhcbM1*, and *ΔLhcbM5* mutants shown in (a). Sample loading was normalized to 3 μg Chl/lane. (c) Absorption spectra of the PSI-LHC(-LHCII) supercomplexes obtained from WT (blue-line), *ΔLhcbM1* (green-line), and *ΔLhcbM5* (red-line) mutants. (d) Fluorescence quenching of the WT strain (blue-line) and ΔLhcbM5 mutant strain (red-line) under state 1 and state 2 conditions, which were induced under far-red (FR-) and blue-light conditions. Data are means ± SE, n = 6 biological replicates. (e) SILAC analysis of the LHC proteins and PsaB (as a control) in the PSI-LHCI-LHCII supercomplexes from the *ΔLhcbM1* mutant compared to that from WT. The relative amounts of each subunit for the PSI-LHCI supercomplex were determined based on the ^14^N:^15^N isotopomer values derived from chromatograms using the stable isotope (^15^N)-labeled LC-MS/MS spectrometry. Each amount was normalized with PsaA abundance. Data shown are the mean ± SE (n = 3). (f) Stromal side view of the structure of *Cr*PSI-LHCI-LHCII supercomplex purified from *ΔLhcbM1* mutant. LhcbM3 that replaces LhcbM1 in the mutant is highlighted in surface mode and colored orange.

Previous data suggested that the absence of LhcbM1 does not affect state transitions ^4,23^, whereas the structure reveals that phosphorylated LhcbM1 does contribute to the formation of state 2 supercomplex in *C. reinhardtii* (Fig. 2b). Curiously, the PSI-LHCI-LHCII supercomplex indeed exists in the *ΔLhcbM1* mutant (lacking LhcbM1, also termed *npq5*), and shows absorption spectrum very similar to those of wild-type supercomplex, although with less amount (Fig. 4a-c). These findings agree well with previous analysis of this mutant strain ^4,23^, and suggest that other LhcbM protein(s) could replace LhcbM1 and participate in forming the state 2 supercomplex in the *ΔLhcbM1* mutant. We then quantified the relative amounts of LhcbM proteins in PSI-LHCI-LHCII supercomplex from *ΔLhcbM1* mutant compared with those in WT cells. After purifying the PSI-LHCI-LHCII supercomplexes, we performed Stable Isotope Labeling with Amino acids in Cell culture (SILAC) analysis to calculate the ^14^N/^15^N ratio of each detected polypeptides corresponding to the ratio of abundance of target proteins in *ΔLhcbM1 versus* in WT cells (Fig. 4e). The results indicate that LhcbM3 abundance is greatly increased in the *ΔLhcbM1* mutant, and the amount of LhcbM2/7 is also higher than that of wild type. On the other hand, all Lhca proteins were not much affected in the PSI-LHCI-LHCII supercomplexes of the *ΔLhcbM1* mutant.

We further solved the cryo-EM structure of PSI-LHCI-LHCII supercomplex from the *ΔLhcbM1* mutant at an overall resolution of 3.16 Å (Fig. 4f, Supplementary Fig. 7). The map clearly shows two LHCII trimers binding at the positions same as their counterparts in the native supercomplex. LHCII-2 is unchanged compared with those in the native supercomplex structure, whereas one Type I isoform replaces LhcbM1 in LHCII-1 trimer. This Type I subunit was assigned as LhcbM3 according to characteristic features of cryo-EM densities (Supplementary Fig. 8) and the SILAC analysis result (Fig. 4e). In the mutant structure, the N-terminal region of LhcbM3 was less built (without the phosphorylation site) and orientates differently compared to that of LhcbM1, while the overall structure of PSI-LHCI-LHCII from the *ΔLhcbM1* mutant highly resembles that of native supercomplex (Supplementary Fig. 9a). The different binding pattern between LhcbM3 and LhcbM1 with the PSI core may lead to the slight pivot of LHCII-1 toward the lumen in the mutant structure, with the outmost region of LhcbM2/7 exhibiting the largest movement of about 2 Å (Supplementary Fig. 9b), whereas the shift at regions of LHCII-1 close to the PSI core is negligible (Supplementary Fig. 9c). Therefore, the results indicate that the absence of LhcbM1 does not affect the formation of PSI-LHCI-LHCII supercomplex, because LhcbM3, with slight adjustment, takes its place in mediating the association of LHCII-1 with PSI.

### PsaO and Lhca9-Lhca2 heterodimer mediating LHCII-PSI assembly

While the PSI-LHCI moiety in the supercomplex shows overall features similar to the previously reported *Cr*PSI-LHCI structures ^25,26^, the PsaO subunit absent in the latter is now observed and modeled at the interface between PSI and LHCII-1 (Fig. 1a). PsaO adopts a folding similar to its homologs from maize and red alga ^19,28^, possessing two TMHs and one amphiphilic helix along the luminal surface, and binding three chlorophyll molecules (Supplementary Fig. 10, Supplementary Table 2). In addition, two β-carotenes were found in *C. reinhardtii* PsaO (absent in plant and red algal PsaO), located at grooves formed by the two TMHs from opposite sides. PsaO is located at the same position in state 2 supercomplex from both plants and green algae, and serves to bridge LHCII-1 and PsaA (Fig. 1b,c).

Compared with the previously reported cryo-EM structures of *Cr*PSI-LHCI complex^25^, the side layer antennae Lhca2 and Lhca9 associate with the core more stably in the state 2 supercomplex as the LHCII-2 trimer serves to stabilize them by shielding the core-Lhca2 interface (Fig. 1a). The well-defined densities (Supplementary Fig. 4) allow the unambiguous modeling of the two LHCIs including their loop regions (Fig. 5a). Both Lhca2 and Lhca9 possess AC loops closely packed against helix C and much shorter than the counterparts of other Lhcas. As a result, they do not bind a chlorophyll at the 608 (nomenclature according to ref ^29^) site conserved in other LHCI proteins, as it would clash with their AC loop (Fig. 5b). Therefore, Lhca2 and Lhca9 contain the least chlorophylls (13 in Lhca2 and 12 in Lhca9) among all the LHCIs in *C. reinhardtii* (Supplementary Table 2). Moreover, Lhca2 and Lhca9 both possess three carotenoids with two located at the conserved L1 and L2 sites crucial for the folding and stability of Lhc proteins ^30,31^. Unlike other Lhcas, they do not bind a carotenoid at the conserved N1 site, but contain a novel carotenoid binding site partially overlapping with the N1 site. Thus, the third carotenoid-binding site in Lhca2 and Lhca9 is named N2 site. The polyene chain of N2-carotenoid is nearly parallel with helix B, and interacts with Chl 604 in both Lhca2 and Lhca9 (Fig. 5b).

**Figure 5.**
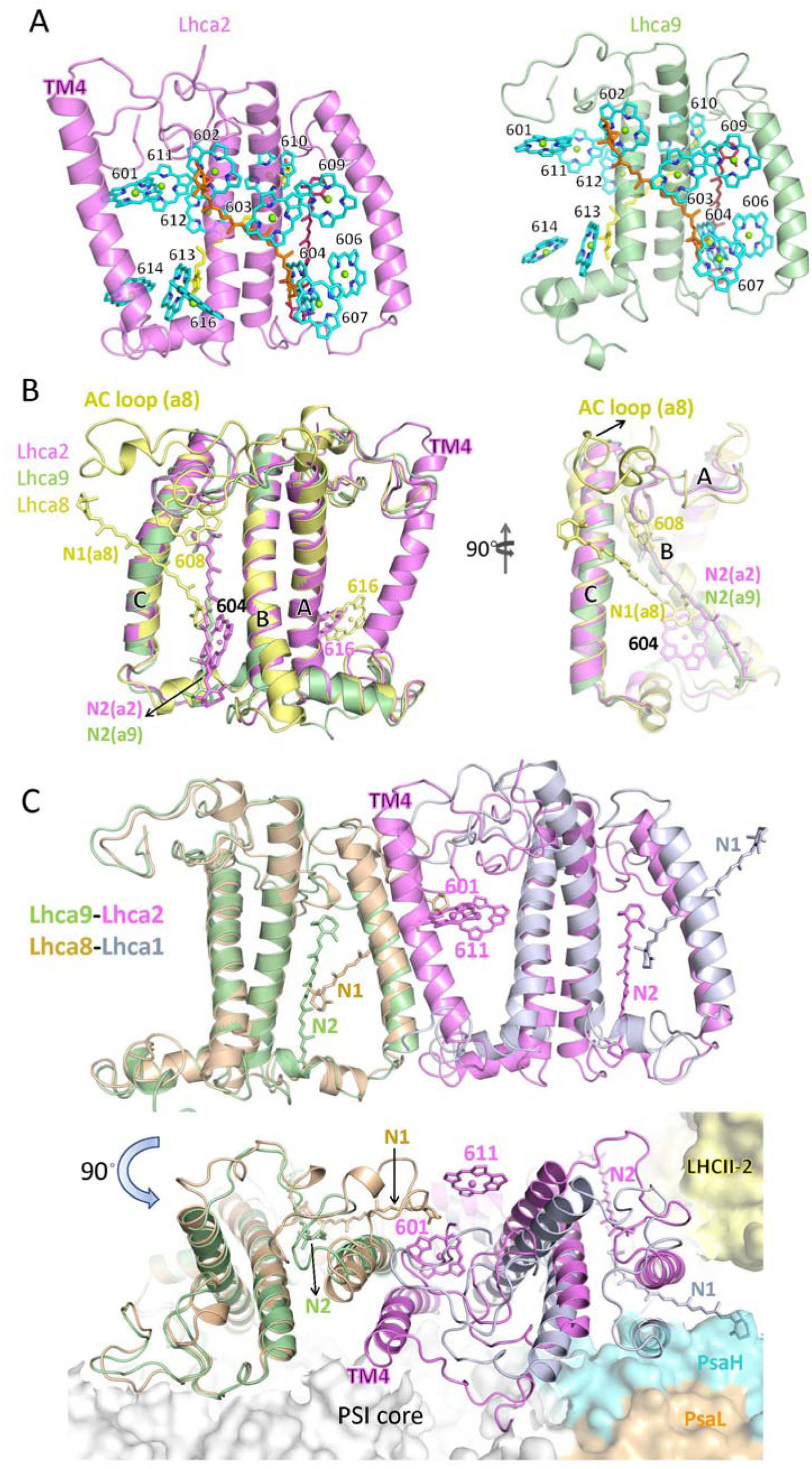
Overall structure of the side layer antennae Lhca2 and Lhca9 from *Cr*PSI-LHCI-LHCII supercomplex. (a) Cartoon representation of *Cr*Lhca2 and *Cr*Lhca9. Chlorophyll are shown in cyan sticks and labeled. Carotenoids at L1, L2 and N2 site are colored yellow, orange and hot pink, respectively. (b) Structural comparison of Lhca2, Lhca9 and Lhca8 (representative of other Lhca proteins in *C. reinhardtii*). Non-conserved pigment molecules and Chl 604 are shown in sticks and labeled. Helices B, C and A are indicated. (c) Structural comparison of Lhca9-Lhca2 heterodimer and Lhca8-Lhca1 heterodimer (representative of other Lhca heterodimers in *C. reinhardtii*) superposed on Lhca9. Pigment molecules involved in the intra-dimer interaction are shown in stick mode and labeled.

The LHCI complexes of PSI usually form heterodimers, with the N1-carotenoid of one LHCI inserting into the space between Chls 601 and 611 from a neighboring LHCI, thus contributing to the dimer stabilization (Fig. 5c). In the side LHCI layer of PSI, Lhca9 does not contain N1-carotenoid. Meanwhile, Lhca2 is different from all other Lhca proteins as it possesses a fourth TMH (Fig. 5a) which is close to and almost parallel with the helix C of Lhca9 (Fig. 5c). Thereby, the fourth TMH of Lhca2 compensates for the loss of N1-carotenoid and mediates the assembly of Lhca9-Lhca2 heterodimer. The distinctively tight dimerization interface may result in the less curved shape of Lhca9-Lhca2 dimer compared with other LHCI heterodimers (Fig. 5c). The flattened Lhca9-Lhca2 heterodimer may accommodate PsaH better at the PsaL-Lhca2 interface and provide a binding site for the association of LHCII-2 trimer. Moreover, binding of LHCII-2 strengthens the association of Lhca9-Lhca2 heterodimer with the core. Therefore, the Lhca2-LhcbM5 interaction is crucial for the mutual stabilization between Lhca9-Lhca2 heterodimer and LHCII-2 trimer in *C. reinhardtii* state 2 supercomplex.

### Pigment arrangement and the potential energy transfer pathways in *Cr*PSI-LHCI-LHCII

The high-resolution cryo-EM maps allowed us to identify most pigment molecules and other cofactors within the supercomplex (Fig. 6, Supplementary Fig. 4). The two LHCII trimers and Lhca9-Lhca2 heterodimer collectively form an antenna belt (Fig. 1a), potentially allowing the excitation energy equilibration among neighboring antenna complexes. In the structure of native *Cr*PSI-LHCI-LHCII supercomplex, excitation energy transfer might occur between LhcbM5 and Lhca2, or between LhcbM5 and LhcbM1. In addition, LhcbM2/7 from LHCII-1 and LhcbM3/M4 from LHCII-2 may also transfer energy to each other. Interestingly, the distances of the interfacial chlorophylls within the belt at the luminal layer are usually shorter than those at the stromal side (Fig. 6), implying that the energy transfer at the luminal side is likely more efficient within the belt.

**Figure 6.**
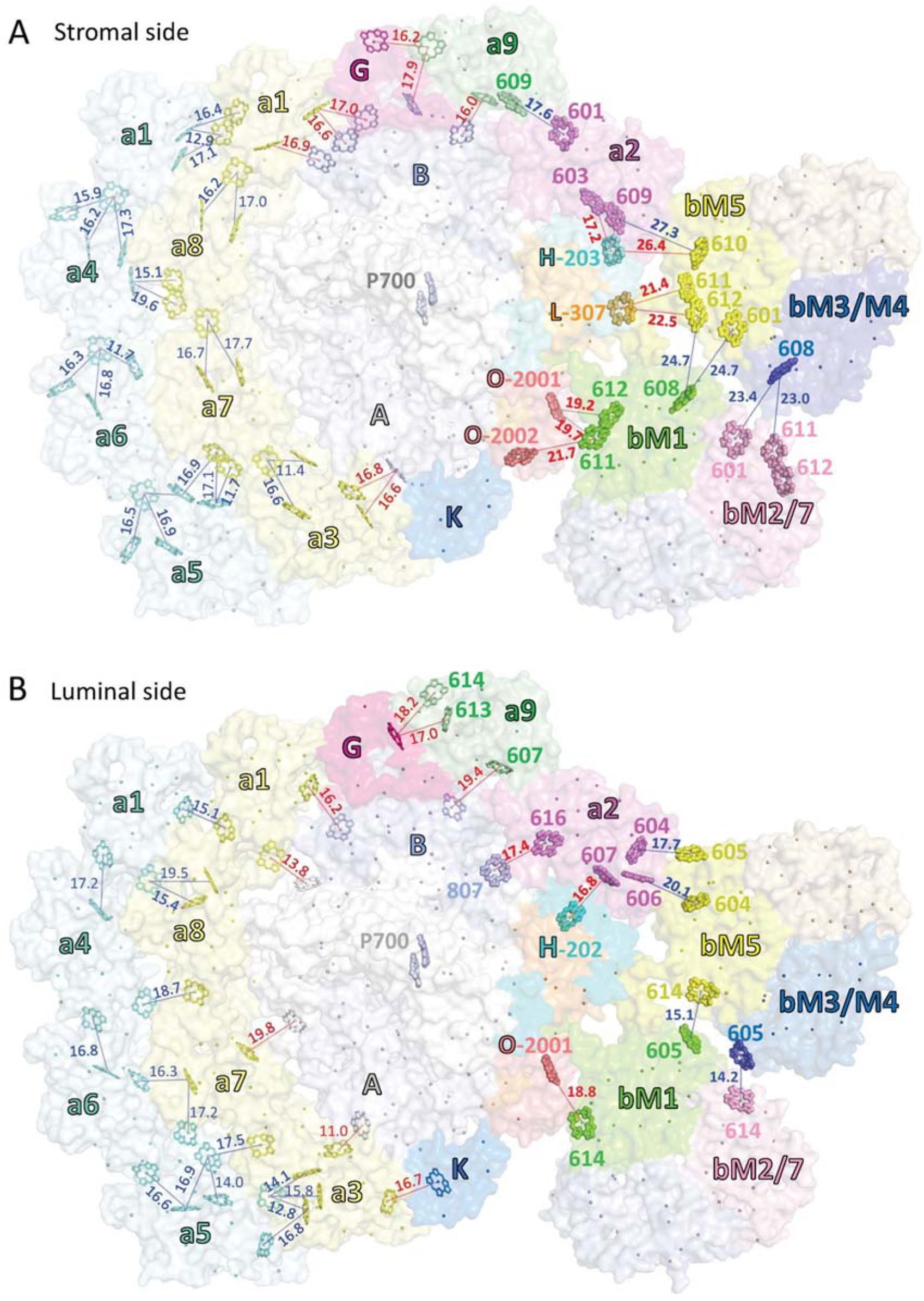
Potential energy transfer pathways within native *Cr*PSI-LHCI-LHCII supercomplex. (a-b) Stromal-side view of the chlorophylls within the *Cr*PSI-LHCI-LHCII supercomplex at the stromal layer (a) and luminal layer (b). Plausible EET pathways between antennas and between antenna and the core are indicated by blue and red lines, with Mg-to-Mg distances (Å) indicated. The chlorophylls at the core-LHCII or Lhca2-LHCII interface and the special pair (P700) are highlighted in ball-and-stick mode. Other chlorophylls are shown as lines or spheres at their central-Mg positions. For clarity, the phytol chains of chlorophylls are omitted.

The chlorophylls located at the interface between PSI and LHCII trimers establish the potential EET pathways from LHCII to PSI. Both LHCII trimers may transfer excitation energy to the PSI core through the Chl 611-612-610 cluster, which is associated with the lowest energy state in plant LHCII ^32^ and might be crucial in relaying EET from LHCII trimers to the core. In LHCII-1, the Chls 611-612 and 614 from LhcbM1 have close distances with Chl 2001 of PsaO, ensuring the efficient EET from LHCII-1 to the core (Fig. 6). In LHCII-2, both Chls 611 and 612 from LhcbM5 are located at distances of approximately 22 Å from Chl 307 of PsaL (Fig. 6), which may constitute the major EET pathways from LHCII-2 to the PSI core. It is apparent that the distances of interfacial Chl pairs between LHCII and PSI core are longer than those at the PSI core-LHCI interface. The observation is consistent with previous data showing that the EET from LHCII to PSI-LHCI is efficient, but it is relatively slow when compared to the process from LHCI to PSI core ^33^.

The PSI-LHCI-LHCII supercomplex from *ΔLhcbM1* mutant has similar pigment arrangement pattern compared to the native supercomplex (Supplementary Fig. 9a), despite that they have slightly shifted LHCII-1 trimer with different protein composition. This structural feature probably ensures that state transition occurs normally in the *ΔLhcbM1* mutant ^4^. However, further analysis is required to clarify the possible difference in the EET between the state 2 supercomplexes in the WT and the *ΔLhcbM1* mutant.

## Discussion

State transitions are a fundamental regulatory process crucial for photosynthetic acclimation in eukaryotic phototrophs. Our *Cr*PSI-LHCI-LHCII structures provide the overall architecture and detailed subunit interactions within the green algal state 2 supercomplex. We showed that two LHCII trimers associate with PSI through three core subunits (PsaO, PsaH and PsaL) and Lhca2 (Fig. 1a). The three core subunits were demonstrated to be crucial for state transitions in plants ^34–36^, and should play similarly pivotal role for state transitions in *C. reinhardtii* based on the similarity of their sequences and structures between the homologs from green algae and plants. Interestingly, PsaO was only observed in state 2 supercomplex in both plants and green algae, while being absent in all previously reported PSI-LHCI structures from green lineage. In contrast, PsaO binds to PsaA stably in red alga which does not have LHCII ^37^. These structural features suggest that PsaO might be loosely associated with PsaA in the green lineage, while binding of LHCII under state 2 conditions stabilizes it at the PsaA-LHCII interface. Besides, PsaO plays a pivotal role in facilitating the assembly and EET between LHCII trimer and PsaA.

The two LHCII trimers in the *Cr*PSI-LHCI-LHCII supercomplex are composed of LhcbM1-LhcbM2/7-LhcbM3/M4 and LhcbM5-LhcbM2/7-LhcbM3/M4. Intriguingly, the LHCII trimers with these particular compositions were identified in the free LHCII fractions in *C. reinhardtii* and shown to exhibit energy dissipation when aggregation was induced under low pH conditions ^38^. The previous result highlighted the quenching capabilities of LhcbM1 and LhcbM5 ^38^, while both isoforms are now proved to be phosphorylated and contribute to the state 2 supercomplex formation (Fig. 2), although with distinct pattern and significance. LhcbM1 is dispensable and could be replaced by LhcbM3 when LhcbM1 is not available (Fig. 4). The capability of LhcbM3 to substitute LhcbM1 could be rationalized by the fact that LhcbM3 and LhcbM1 have similar N-terminal tail length and pThr position ^24^. However, we did not observe cryo-EM density corresponding to the phosphorylated N-terminal tail of LhcbM3 at the surface pocket of PSI where the LhcbM1 N-terminal tail binds, indicating that the phosphorylated N-terminal region of LhcbM3 is mobile and its binding site on PSI surface may be shifted slightly from that of LhcbM1. The pocket holding the N-terminal tail of LhcbM1 is highly acidic (Fig. 2d), which should be favorable for accommodating the two basic residues preceding pThr27 in LhcbM1, but may not be ideal for harboring the less basic N-terminal tail of LhcbM3 (^25^GKtAA^29^ in LhcbM3 *versus* ^25^RRtVK^29^ in LhcbM1).

In contrast to LhcbM1, our results demonstrated that LhcbM5 is essential for state transitions in *C. reinhardtii* (Fig. 4). A recent report showed that only cells from *pph1;pbcp* double mutant are locked in state 2, and both phosphatases PPH1 and PBCP are able to dephosphorylate LhcbM5 ^13^. This finding together with our structural observation suggested that the phosphorylated N-terminal region of LhcbM5 is prerequisite for state 2 supercomplex formation. LhcbM5 is unique as it has never been detected in the PSII-LHCII complex ^39,40^, but only present in the PSI-LHCI-LHCII supercomplex under state 2 conditions ^21,27^. It is possible that LhcbM5 does not bind to PSII core, and LHCII-2 trimer containing LhcbM5 may only exist in the LHCII pool region. Under state 2 conditions, phosphorylation of the N-terminal tail of LhcbM5 enhances its affinity with the PSI-LHCI, leading to the association of a portion of LHCII-2 trimers with the PSI core, which further stabilizes the binding site of LHCII-1. Therefore, we propose that the LHCII-2 trimer is not shuttled from PSII-LHCII complex, but could be delivered from the bulk LHCII pool in the thylakoid membranes, as it may be more efficient for state transitions to occur. Moreover, the remaining LHCII-2 together with the mobile LHCII-1 trimers may aggregate and turn to quenched state as previously suggested ^38^.

The state 2 supercomplex in plants is smaller than the one from *C. reinhardtii* and only contains one LHCII trimer. Interestingly, only the LHCII-1 binding site is kept in plants during evolution (Fig. 1b), and Lhcb2 (with the phosphorylated N-terminal motif identical to that of LhcbM1 in green algae) becomes the crucial one for the formation of state 2 supercomplex in plants. As higher plants face stronger and more fluctuating light conditions upon transition from aqueous to terrestrial environments, they have lost several Lhca proteins, including the side layer antennae (Lhca9 and Lhca2) which provide a docking site for LHCII-2. Consequently, the LHCII-2 binding site is concomitantly lost in plants, and only the binding site for LHCII-1 has been preserved.

## Method

### Algal strains

*Chlamydomonas reinhardtii* control strain 137c (wild type; WT) was obtained from the Chlamydomonas Center (https://www.chlamycollection.org/). The *pph1;pbcp* double mutant strain is a kind gift from Professor Michel Goldschmidt-Clermont at University of Geneva ^13^. The *npq5* (*ΔLhcbM1*) strain was generated in a previous study ^4^. The *ΔLhcbM5* strain was obtained during the genetic screening in the previous study ^41^. In brief, we performed insertional mutagenesis using *aph7* (hygromycin resistant) cassette using the *LHCSR1–Luc717* strain harboring a reporter construct expressing a LHCSR1-Luciferase fusion ^41^. The exact *aph7* insertion site in the *ΔLhcbM5* strain was confirmed to be in the sixth intron of the *LhcbM5* gene by sequence analysis (Fig. S6A). Furthermore, we confirmed that LhcbM5 polypeptide was not detectable in the *ΔLhcbM5* strain by immunoblotting analysis using an LhcbM5 antibody (Fig. S6B); the antibody was raised against LhcbM5 peptide RVNGGPAGEGLDK.

### Growth conditions

State 2 in *C. reinhardtii* could be induced by either incubating the cells under the anaerobic condition in the dark ^42–44^ or illuminating the cells with high light ^45^. For *C. reinhardtii* strain *137c* (WT), *ΔLhcbM1* and *ΔLhcbM5*, cells were grown in Tris-acetate-phosphate (TAP) medium ^46^ under dim light (< 20 μE m^−2^ s^−1^) at 23°C until they reached the mid-log growth phase. The culture medium was exchanged with modified high-salt (HS) minimal medium ^41^. After the medium exchange with modified HS minimal medium, the cells were grown photoautotrophically under continuous 100 μmol/m^2^/s white fluorescent light with 5% CO_2_ bubbling at 23°C. For the stoichiometry analysis, the WT cells were grown in the modified HS minimal medium using ^15^N-labeled NH_4_Cl with 5% CO_2_ bubbling at 23°C under continuous 100 μmol/m^2^/s white fluorescent light.

For *C. reinhardtii pph1;pbcp* double mutant strain, cells were grown in Tris-acetate-phosphate medium ^46^ at 23°C under low light (20 μmol m^−2^ s^−1^ photons) conditions, and were treated with high light condition (330 μmol m^−2^ s^−1^ photons) for one hour before harvesting.

### Isolation of PSI-LHCI-LHCII supercomplexes from WT, *ΔLhcbM1* and *ΔLhcbM5* strains

Prior to the thylakoid membrane isolation, the cells were treated with carbonylcyanide-3-chlorophenylhydrazone (CCCP) at a final concentration 5 µM for 15 min in the dark. This process leads the cells into state 2 ^47^. Thylakoid membranes from the WT, *ΔLhcbM1*, and *ΔLhcbM5* strains after state 2 induction were isolated as previously reported ^6^. To purify the photosynthetic supercomplexes from the isolated thylakoids, we used amphipol A8-35 as previously reported ^48^. In brief, thylakoid membranes were detergent-solubilized as described in a previous report ^6^, then immediately mixed with A8-35 at a final concentration of 1.0 % and incubated on ice for 15 minutes. The A8-35-treated membranes were separated by detergent-free sucrose density gradient ultracentrifugation at 230,000 *g* for 16 hours. The bands corresponding to PSI-LHCI and PSI-LHCI-LHCII samples (Fig. 4A) were obtained and used for biochemical analysis.

### Isolation of PSI-LHCI-LHCII supercomplex from *pph1;pbcp* mutant strain

The thylakoid membranes were prepared following the previously published protocol^49^. In brief, collected cells were resuspended in Lysis Buffer (containing 25 mM HEPES, 0.3 M Sucrose, 5 mM MgCl_2_), then passed through a high-pressure homogenizer device (AH-2010, ATS Engineering) at a pressure of 500 bar for 3 times. The homogenate was centrifuged, and the thylakoid containing pellet was washed with washing buffer (5 mM HEPES, 0.3 M Sucrose, 10 mM EDTA) once. The thylakoid membranes were further separated from contaminants through a discontinuous sucrose density gradient containing three layers of 0.5M/1.3M/1.8M sucrose respectively ^49^. The thylakoid membrane sample at the interface between two layers was harvested and further solubilized with 1.0% (w/V) dodecyl-α-D-maltopyranoside (α-DDM) at 4°C for 10 min at a chlorophyll concentration of 0.4 mg/ml. The insoluble fractions were discarded after centrifugation at 10,000 g for 5 min. Photosynthetic supercomplexes were separated through sucrose density gradient (SDG) ultra-centrifugation at 120,000 *g* for 18 h, with the matrix containing 5-35% sucrose, 25 mM MES pH 6.5 and 0.02% α-DDM. Bands corresponding to PSI-LHCI and PSI-LHCI-LHCII supercomplexes were collected for further study (Fig. S1A).

### Characterization of *Cr*PSI-LHCI-LHCII supercomplexes

For each SDG band from WT, *ΔLhcbM1* and *ΔLhcbM5* strains, the intactness of the PSI-LHCI(-LHCII) supercomplexes were analyzed using 3-12% Native-Gel Bis-Tris NativePAGE^TM^ gel (Fig. 4B) following the manufacturer’s instructions (Thermo Fisher Scientific). The identities of proteins were further verified by mass spectroscopy analysis of the in-gel trypsin-digestion products of the excised CN–PAGE bands. Absorption spectrum analysis was carried out by using an IMPLEN NP-80 UV-Vis spectrophotometer, with a wavelength range of 350-750 nm (Fig. 4C).

For each SDG band from *pph1;pbcp* double mutant strain, the protein composition was analyzed using 10–15% gradient sodium dodecyl sulfate–polyacrylamide gel electrophoresis (SDS–PAGE) (Fig. S1B). The gels were stained with Coomassie Brilliant Blue R250. The identities of proteins were further verified by mass spectroscopy analysis of the in-gel protease-digestion products of the excised SDS–PAGE bands. Absorption spectrum analysis was carried out by using a TECAN 96-well plate reader, with a wavelength range of 400-750 nm (Fig. S1C). High performance liquid chromatography (HPLC) was performed as previously described ^50^, following the protocol reported by Färber et al ^51^ (Fig. S1D).

### Stoichiometry analysis

Stoichiometry analysis was performed as described previously ^52^. Stable isotope (^15^N) labeling by amino acids in cell culture (SILAC) method was conducted prior to the LC-MS/MS analysis. From the ^14^N:^15^N isotopomer values derived from the LC-MS/MS chromatograms, the WT: *ΔLhcbM1* ratios were calculated for each PSI-LHCI-LHCII subunit (Fig. 4E).

### Chlorophyll fluorescence quenching analysis

Chlorophyll fluorescence quenching analysis was performed as described previously^53^. The state-transition induction light was provided by monochromatic LED light. Far-red light (720 nm, 12.6 μmol/m^2^/s) and blue light (465 nm, 12.5 μmol/m^2^/s), which preferentially stimulate PSI and PSII, were prepared during the chlorophyll fluorescence measurements using handmade spectroscopy ^47^. All processes were performed at 23°C.

### Grid preparation and data acquisition

For the PSI-LHCI-LHCII samples isolated from WT and *ΔLhcbM1* mutant, before cryo-EM sample preparation, the sucrose was removed by gel-filtration using PD-10 column (GE healthcare) with a sucrose-free buffer (25 mM MES pH 6.5 and 0.5 M betain) and then concentrated in a 100 kDa molecular weight cut-off centrifugal filter unit (Amicon Ultra-15, Merck Millipore). Subsequently, 2.5 μl of the sample (3 mg ml^−1^ Chl) was applied onto R1.2/1.3 Cu 300 mesh holey grids (Quantifoil Micro Tools), which were pretreated with glow discharge using a plasma ion bombarder (PIB-10, Vacuum Device) for 30 s. The grid was blotted for 3-5 s with a force level of 7 at 23 °C and 95% humidity and flash-frozen in liquid ethane using a Vitrobot Mark IV (Thermo Fisher Scientific). The frozen grids were stored in liquid nitrogen.

For the PSI-LHCI-LHCII samples isolated from the *pph1*;*pbcp* strain, before cryo-EM sample preparation, the sucrose was removed by diluting the sample 1000 times with a sucrose-free buffer (25 mM MES pH 6.5 and 0.02% α-DDM) and then concentrated in a 100 kDa molecular weight cut-off centrifugal filter unit (Amicon Ultra-15, Merck Millipore). The dilution and concentration process was repeated 3 times. Subsequently, 3 μl of the sample with a concentration of 5 mg ml^−1^ Chl was loaded onto a H_2_/O_2_ glow-discharged holey carbon grid (Quantifoil 300 mesh, R1.2/1.3). The grid was blotted for 4 s with a force level of 2 at 22 °C and a humidity of 100%, flash-frozen in liquid ethane and transferred to liquid nitrogen for storage.

For structure determination, the micrographs of PSI-LHCI-LHCII supercomplexes from WT, *pph1;pbcp* mutant and *ΔLhcbM1* mutant were collected using SerialEM software suite ^54^, on a 200 kV Talos Arctica electron microscope (Thermo Fisher Scientific) equipped with a Gatan K2 Summit direct detector camera and a GIF quantum energy filter (20 eV). Total of 2725, 4988 and 8259 images were collected for PSI-LHCI-LHCII supercomplexes from WT, *pph1*:*pbcp* mutant and *ΔLhcbM1* mutant strains, using a defocus range between 1.5 μm and 2.2 μm at 130,000 magnification. Images were recorded by beam-image shift data collection methods ^55^. The micrographs were exposed for 10 s and dose fractionated into 32 frames, leading to a total dose of 50 e^−^·Å^−2^.

### Cryo-EM data processing, classification and reconstruction

For the data processing, the 32 frames in each movie stack were aligned, dose weighted and summed using MotionCor2 ^56^, and binned to a pixel size of 1.0 Å. The contrast transfer function (CTF) parameters of summed images were estimated using CTFFIND4.1 ^57^. Further image processing steps were performed in Relion v.3.1 or Relion v3.0 ^58^, including reference-based particle autopicking, 2D/3D classifications, 3D auto-refinement, CTF correction and Bayesian particle polishing. Local maps were generated in UCSF Chimera ^59^, and were further processed into masks using Relion. The detailed data processing workflow was shown in Figs. S2, S3 and S7. The final resolutions of density maps of *Cr*PSI-LHCI-LHCII from *pph1*;*pbcp* and *ΔLhcbM1* mutant strains are 2.84 and 3.16 Å, respectively, estimated based on the gold-standard Fourier shell correlation with 0.143 criterion. To further improve the density of two LHCIIs, focused refinement was used with a soft edged mask around the LHCII-1 and LHCII-2 regions, respectively. The local resolution of the final map was calculated using ResMap ^60^ (Figs. S3, S7).

### Model building and refinement

To construct the structural model of the PSI-LHCI-LHCII supercomplexes from *pph1*;*pbcp* mutant strain, the cryo-EM structures of *Cr*PSI-LHCI complex (PDB code: 6IJO) and the LHCII trimer from *Cr*PSII-LHCII complex (PDB code: 6KAD) were docked into the 2.84 Å resolution map and the focused refinement maps, respectively, using UCSF Chimera, and used as the initial models for further rebuilding and adjustment. The fourth helix of Lhca2 was built manually on the basis of the density map using Coot ^61^. The structure of maize PsaO from the plant state 2 complex (PDB code: 5ZJI) was docked into the map, and the amino acid residues were mutated to the sequences of PsaO from *C. reinhardtii*. To construct the structural model of the PSI-LHCI-LHCII supercomplex from *ΔLhcbM1* mutant strain, the structure of PSI-LHCI-LHCII from *pph1*;*pbcp* mutant was docked into the 3.16 Å resolution map. The LhcbM3 which replaces LhcbM1 was mutated manually according to the sequence. The models were rebuilt and adjusted manually using Coot ^61^. After model building, real-space refinements were performed using Phenix-v1.15^62^, and the geometric restraints of the cofactors and the Chl–ligand relationships were supplied during the refinement. Automatic real-space refinements followed by manual correction in Coot were carried out interactively. The geometries of the final structures were assessed using MolProbity ^63^. A summary of the statistics for data collection and structure refinement of PSI-LHCI-LHCII supercomplexes from *pph1*;*pbcp* mutant and *ΔLhcbM1* mutant strains is provided in Table S1. High-resolution images for publication were prepared using Chimera and PyMOL (Molecular Graphics System, LLC).

### Identification of each LhcbM protein

In the structure of PSI-LHCI-LHCII from *pph1*;*pbcp* mutant, the three subunits in LHCII-1 were assigned as LhcbM1, LhcbM2/7 and LhcbM3/M4, according to specific density features (Fig. S5). The map features in the N-terminal region of LhcbM1 and its specific residue F106 were used to identify LhcbM1 (Fig. S5A). Moreover, LhcbM1 possesses the identical sequence with Lhcb2 for five residues (RRtVK) at the N-terminal tail, which could be superposed well, further confirming the assignment of LhcbM1.

LhcbM5 possess the longest N-terminal region among all LhcbM proteins in *C. reinhardtii*, which shows clear densities in the map. The assignment of LhcbM5 was further verified by the densities of specific residues F197 and F254 (Fig. S5D).

For the identification of LhcbM2/7 in both LHCII trimers, the map features of W32 and T98 exclude the possibility of other LhcbM proteins (Figs. S5C, S5E), as LhcbM3/4/5/6/8/9 contain a phenylalanine in the corresponding position of W32, and LhcbM1 possesses Phe in the corresponding position of T98. The densities of other characteristic sites, including residues F147, P171 and H173 were used to further verify the assignment of LhcbM2/7. LhcbM2 and LhcbM7 could not be distinguished because they share the identical primary sequences for their mature proteins ^64^.

For the identification of LhcbM3/M4 in LHCII-1 and LHCII-2 (Figs. S5B, S5F), the map features of F40 and W48 in the N-terminal regions exclude the possibility of LhcbM1 and LhcbM2/7 (Trp and Phe in the corresponding positions). In addition, the map features of F117 as well as Y52, P179 and Y181 further verify the assignment of Type I isoform. Furthermore, we compared the sequences of five Type I isoforms, and checked the map features of their variant residues. Our analysis suggested that this isoform in both LHCII trimers are LhcbM3 or LhcbM4, as the densities of M213 exclude LhcbM6; M216 exclude LhcbM9; T242 exclude LhcbM8 and LhcbM9 (Figs. S5B, S5F).

In the structure of PSI-LHCI-LHCII from *ΔLhcbM1* mutant, a type I isoform (LhcbM3/4/6/8/9) was identified to substitute LhcbM1, as it fits well to the map of some typical residues (Fig. S8). For example, the map features of W48 and Y52 in the N-terminal regions exclude the possibility of Type III / Type IV (Phe in the corresponding position of W48) and Type II (Leu in the corresponding position of Y52). The map features of Y181 further verify the assignment of Type I isoform. The resolution is not high enough to distinguish among the five type I LhcbM proteins, however, our SILAC result (Fig. 4E) clearly showed that the amount of LhcbM3 is nearly doubled in the *ΔLhcbM1* mutant compared with the WT, while other type I LhcbM proteins show similar amount in both WT and mutant strain. Therefore this LhcbM protein was confidently assigned as LhcbM3 in the structure of PSI-LHCI-LHCII from *ΔLhcbM1* mutant.

## Data availability

The atomic coordinates of the *Cr*PSI-LHCI-LHCII supercomplex has been deposited in the Protein Data Bank with accession code #### (native supercomplex from *pph1;pbcp* mutant) and #### (supercomplex from *ΔLhcbM1* mutant). The cryo-EM map of the supercomplex has been deposited in the Electron Microscopy Data Bank with accession code EMD-#### and EMD-####.

## Acknowledgements

We thank Prof. Michel Goldschmidt-Clermont at the University of Geneva for the kind gift of *pph1;pbcp* strain and insightful discussion; L. H. Chen, X. J. Huang, B. L. Zhu and F. Sun at the Center for Biological Imaging (IBP, CAS) for the support in cryo-EM data collection; X. Sheng, D. F. Song and X. Z. Zhang for discussion on cryo-EM data processing methods; X. B. Liang for the assistance in sample preparation, data collection and storage. We are grateful to Mrs. Tomoko Mori and Ms. Yumiko Makino for providing technical assistance with LC-MS/MS analysis. We also appreciate Mr. Akimasa Watanabe for technical assistance and Dr. Eunchul Kim for valuable discussion. The project is funded by the National Key R&D Program of China (2017YFA0503702), the Strategic Priority Research Program of CAS (XDB27020106 and XDB37020101), the National Natural Science Foundation of China (31925024 and 31930064), the Basic Frontier Science Research Program of CAS (ZDBS-LY-SM003) and Grant-in-Aid from Japan Society for the Promotion of Science (JSPS) KAKENHI (16H06553 and JP15H05599). This work was also supported by Functional Genomics Facility, NIBB Core Research Facilities, and by Model Plant Research Facility, NIBB Bioresource Center, and the Cooperative Study Program of National Institute for Physiological Sciences (NIPS).

## Author contributions

R.T., Z.L., J.M. and M.L. conceived and coordinated the project. X.P. and A.L. collected and processed the cryo-EM data. X.P. built and refined the structural models. R.T. performed the biochemical and spectroscopic characterization of the PSI-LHCI(-LHCII) supercomplexes from WT and LhcbM mutants. A.L. prepared the cryo-EM grids for the supercomplex sample from *pph1;pbcp* mutant. C.S. and K.M. prepared the cryo-EM grids for the supercomplex samples from WT and *ΔLhcbM1* mutant. T.Y. generated the *ΔLhcbM5* mutant. K.T. performed qT quenching measurement. X.P., R.T., A.L., Z.L., J.M. and M.L. analyzed the data and wrote the manuscript; all authors discussed and commented on the results and the manuscript.

## Supplementary Materials

**Fig. S1.**
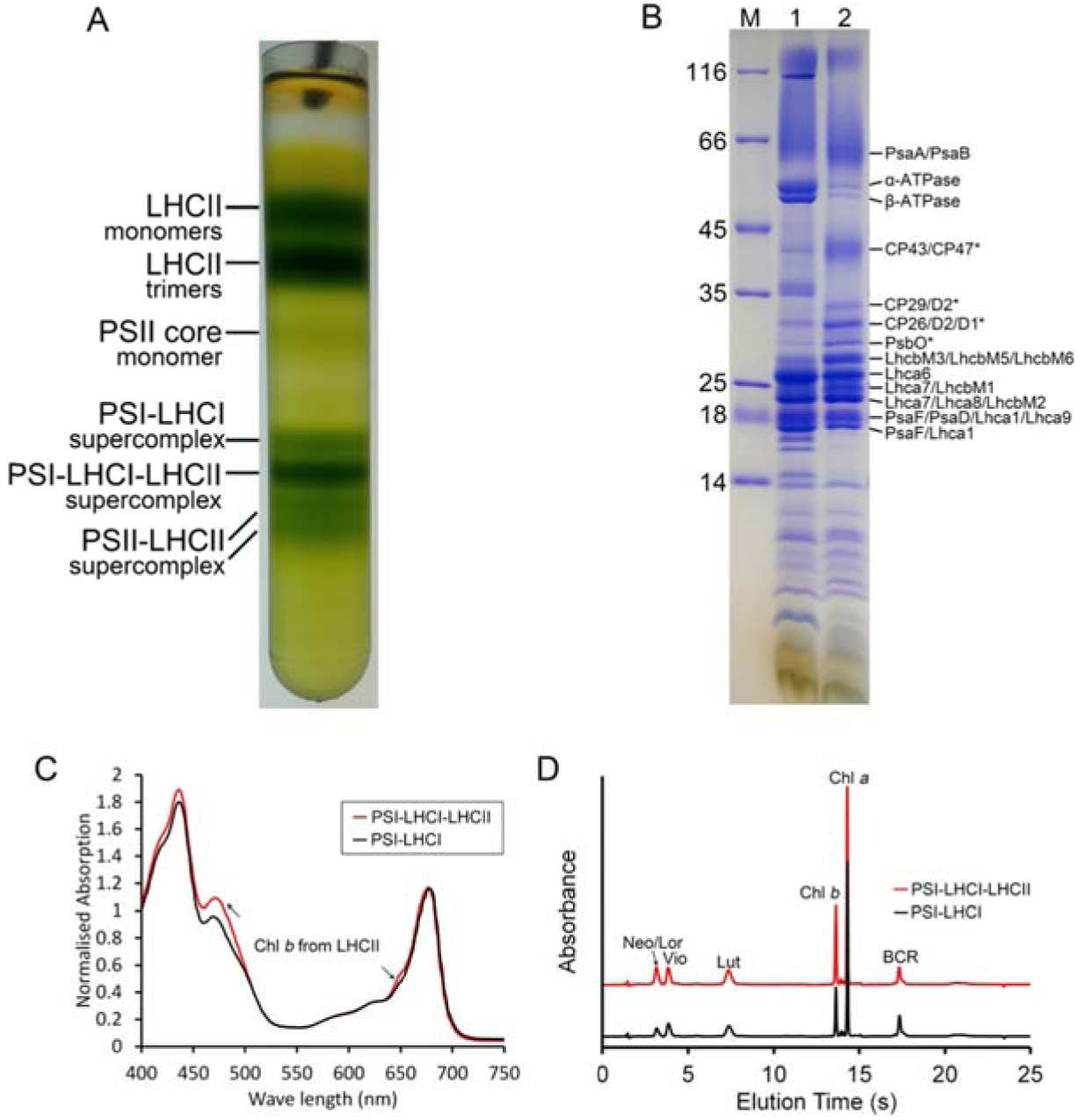
Sample preparation and characterization of native *Cr*PSI-LHCI-LHCII supercomplex from the *pph1;pbcp* mutant strain. (A) Sucrose density gradient of solubilized thylakoid membranes. Each band shown is labeled. (B) SDS-PAGE analysis of thylakoids (Lane 1) and the purified PSI-LHCI-LHCII supercomplex (Lane 2). The protein composition of each Coomassie band was indicated based on the mass spectrometry and proteomics data analysis. The bands corresponding to PSII subunits are indicated by asterisk. (C) Room-temperature absorption spectra of PSI-LHCI and PSI-LHCI-LHCII samples. The PSI-LHCI-LHCII sample showed higher peaks around 470 and 650 nm (indicated by arrows), demonstrating that the Chl b (from LHCII) content of this fraction is higher than that of PSI-LHCI complex. The spectra were normalized to the maximum in the red region. (D) HPLC analysis of pigment content in PSI-LHCI and PSI-LHCI-LHCII samples. Based on the characteristic absorption spectrum of each peak fraction, the six major pigment peaks separated from the sample are identified as loroxanthin/neoxanthin (Lor/Neo), violaxanthin (Vio), lutein (Lut), chlorophyll *b* (Chl *b*), chlorophyll *a* (Chl *a*) and β-carotene (BCR).

**Fig. S2.**
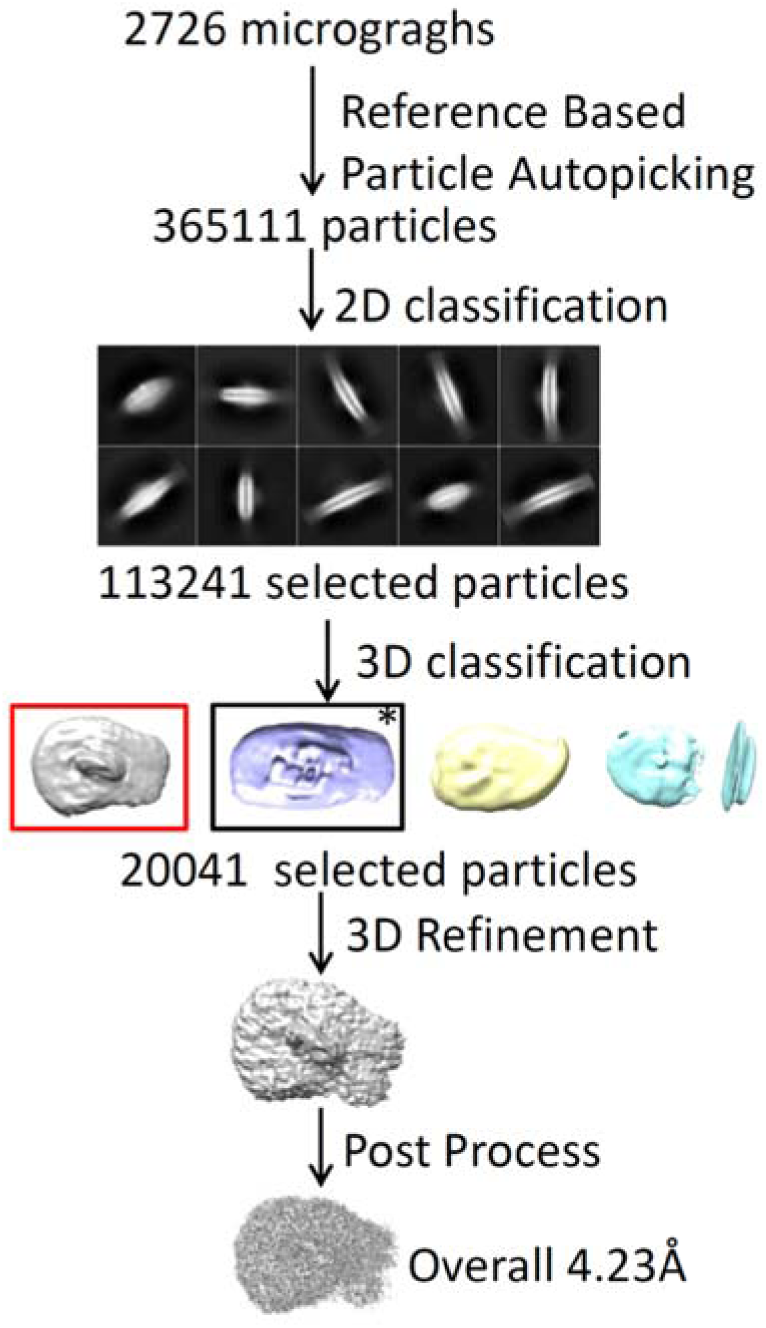
Single particle cryo-EM data processing of *Cr*PSI-LHCI-LHCII supercomplex from *C. reinhardtii* wild type strain. Single particle cryo-EM data processing procedure. The class corresponding to PSII-LHCII complex is framed by black box and indicated by asterisk.

**Fig. S3.**
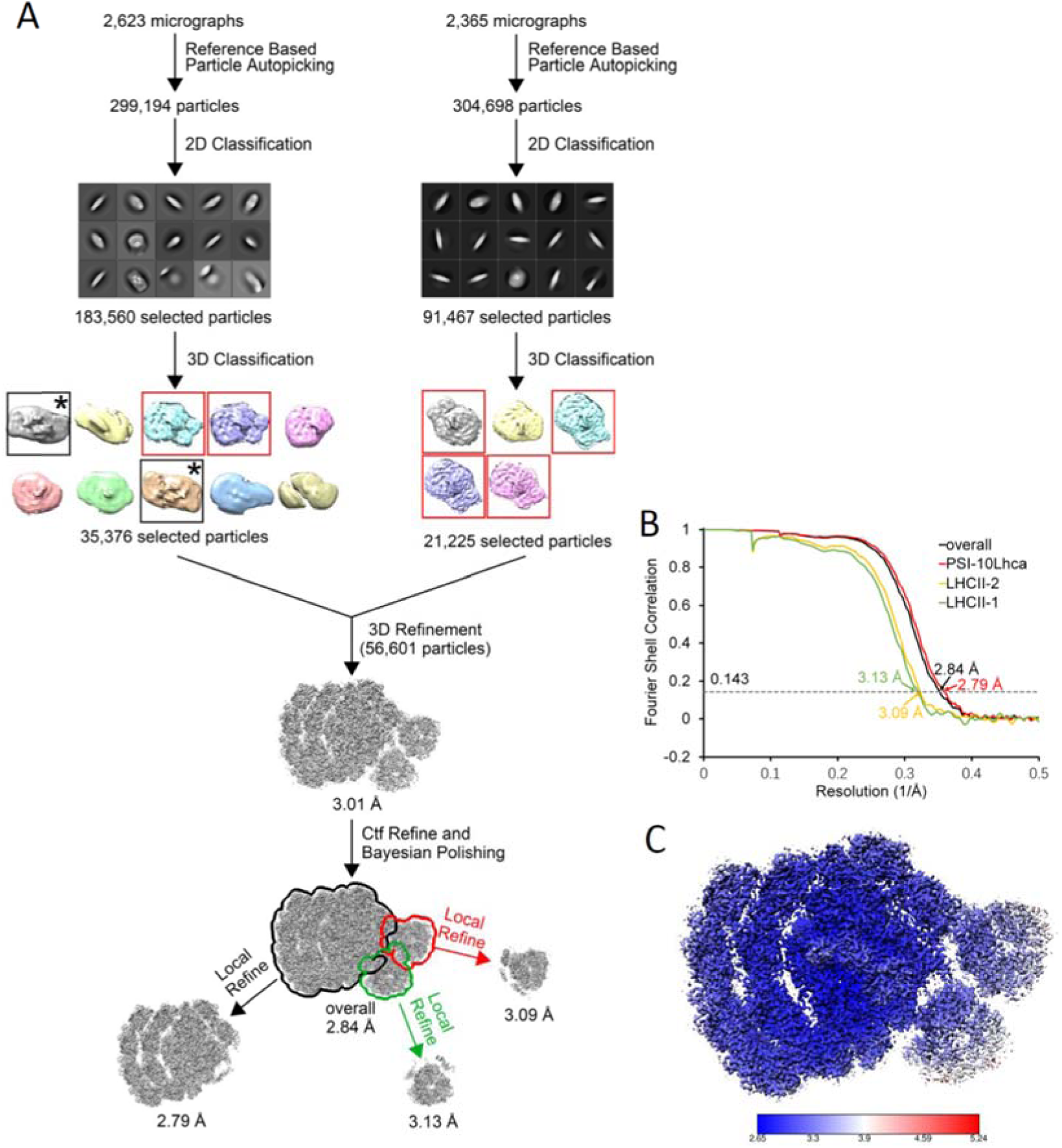
Single particle cryo-EM analysis and evaluation of *Cr*PSI-LHCI-LHCII supercomplex from *pph1;pbcp* mutant strain. (A) Single particle cryo-EM data processing procedure. The class corresponding to PSII-LHCII complex is framed by black box and indicated by asterisk. (B) The gold standard Fourier shell correlation (FSC) curves of the final density map with criterion of 0.143. (C) Local resolution of the cryo-EM map estimated by ResMap.

**Fig. S4.**
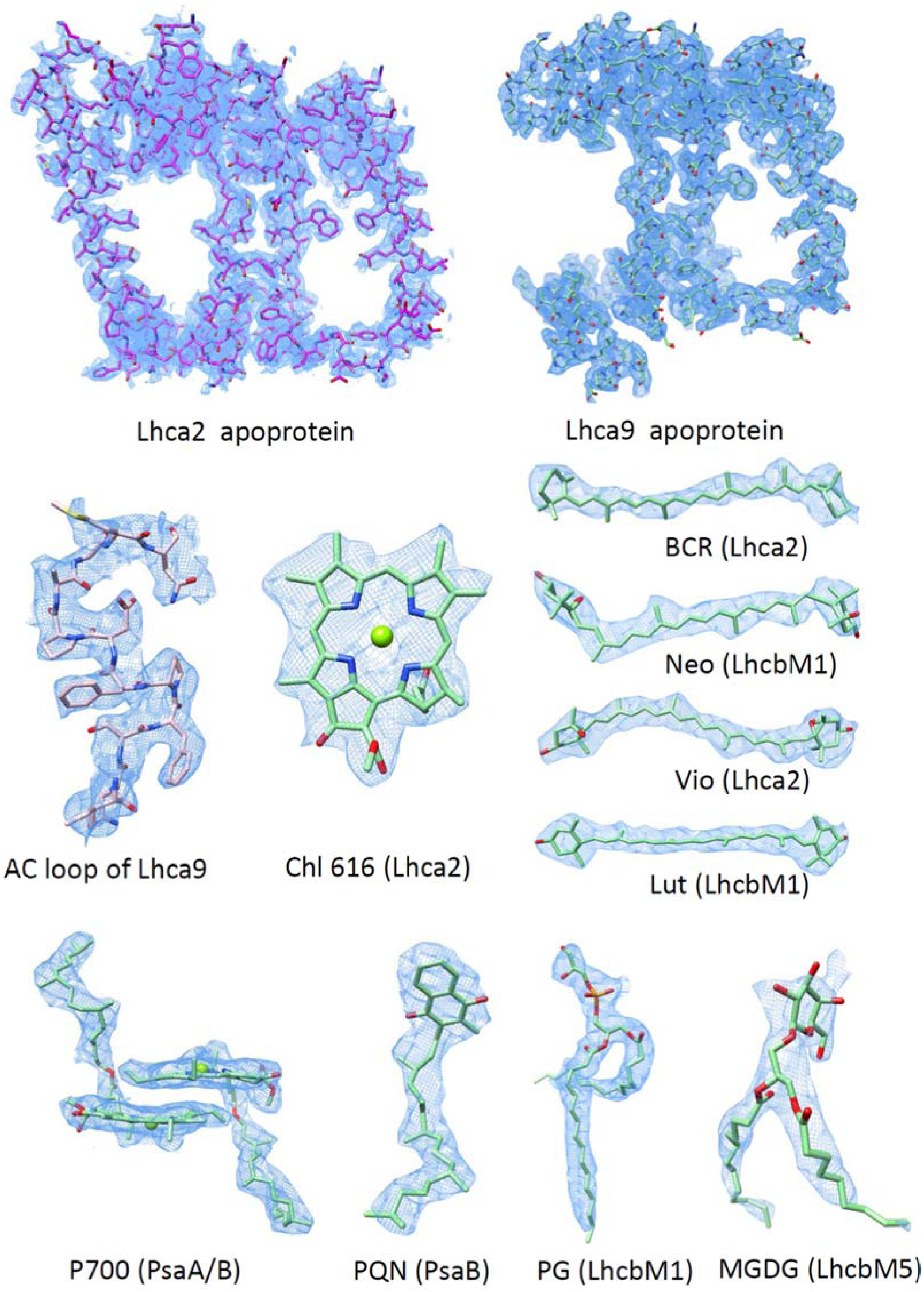
Representative densities of protein subunits and cofactors in the *Cr*PSI-LHCI-LHCII supercomplex structure.

**Fig. S5.**
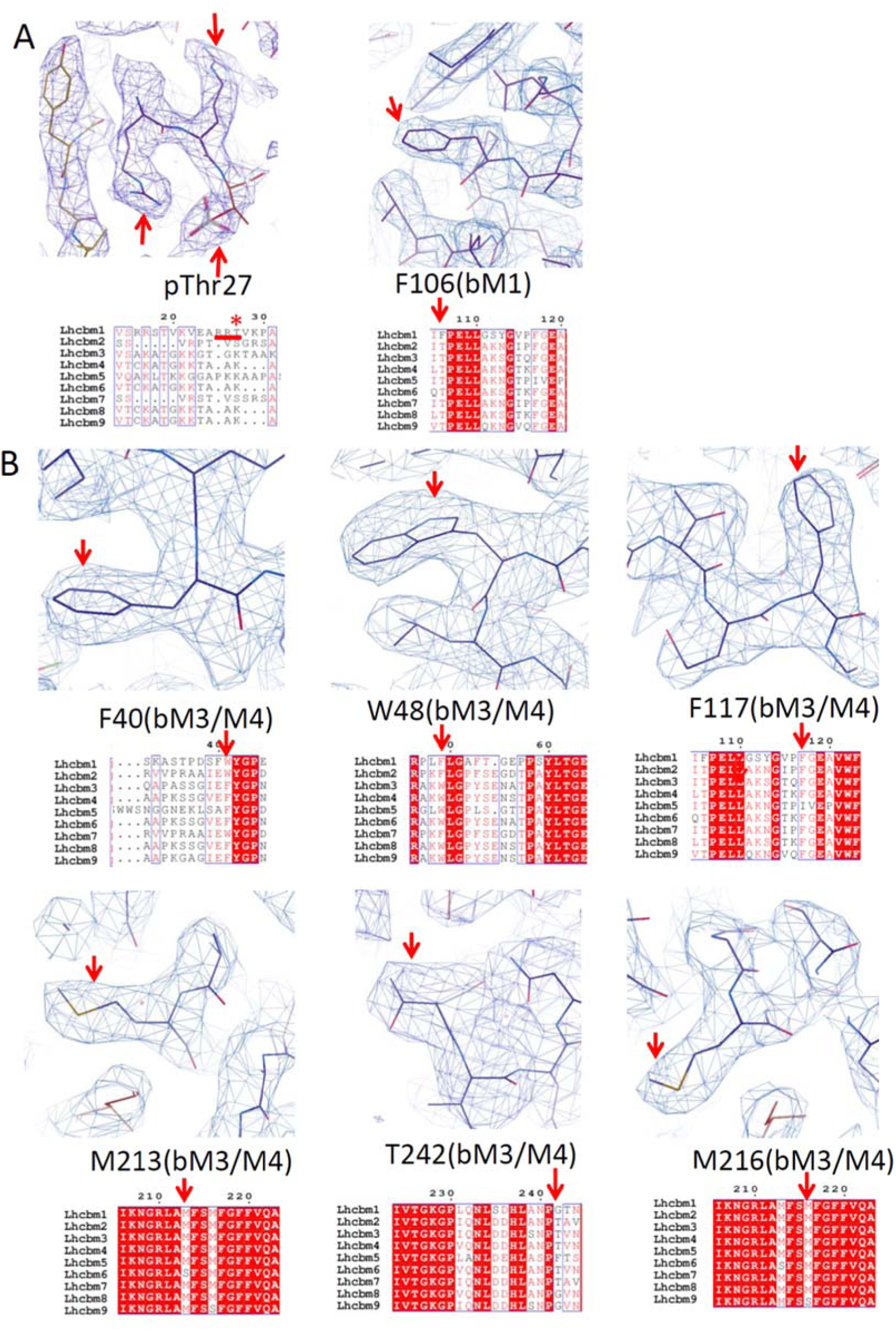
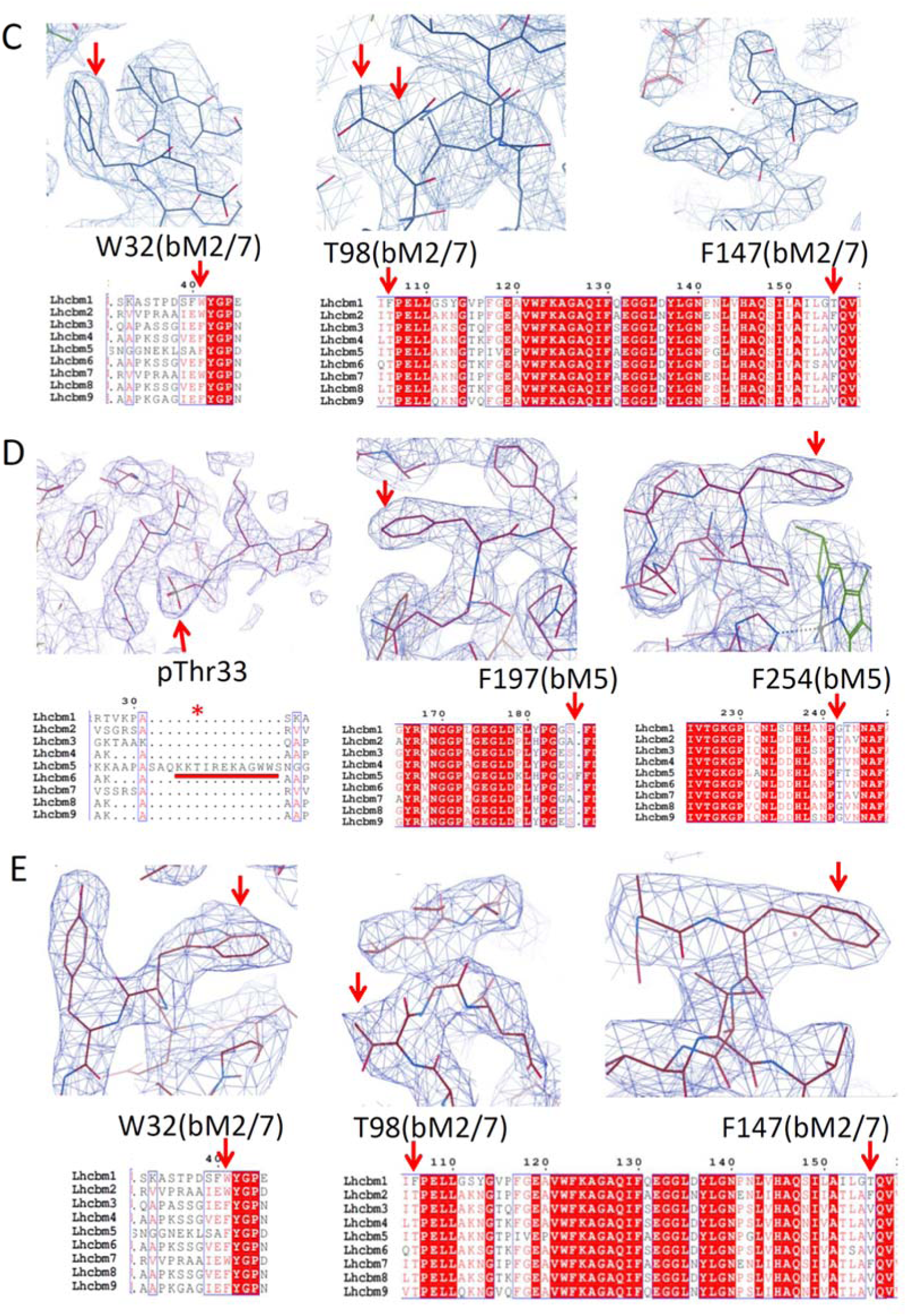
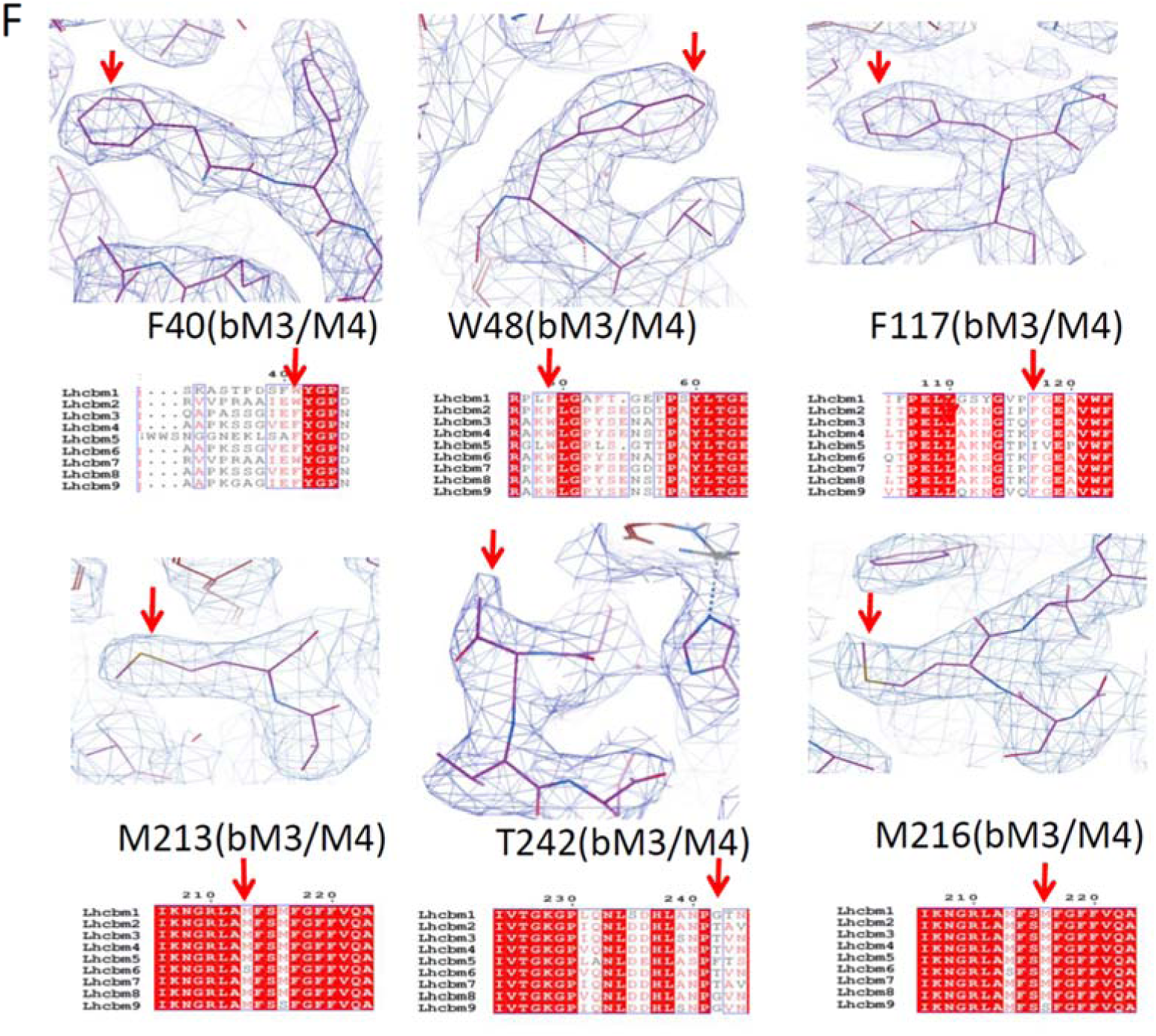
Identification of each LhcbM protein in the *Cr*PSI-LHCI-LHCII supercomplex structure. (A-F) Map features of characteristic residues and the corresponding sequence of LhcbM1 (A), LhcbM3/M4 (B) and LhcbM2/7 (C) from LHCII-1, LhcbM5 (D), LhcbM2/7 (E) and LhcbM3/M4 (F) from LHCII-2. Unique residues used for identification are indicated by red arrows. (A) LhcbM1 possesses unique N-terminal residues (RRt, underlined by a red line) with well-defined density. Moreover, Phe106 shows clear density for its side-chain, which excludes the possibility of all other LhcbM proteins as they have a Thr at the corresponding position. (B) Type I LhcbM proteins have Phe40 and Trp48 in the N-terminal regions, whereas Type III and Type IV have Trp and Phe in the corresponding positions, therefore can be excluded. In addition, Phe117 shows clear density for its side-chain, which excludes the possibility of Type II LhcbM protein as LhcbM5 has a Ile at the corresponding position. The well-defined densities of Met213, Met216 and Thr242 further exclude LhcbM6, LhcbM9, and LhcbM8 and LhcbM9. (C) LhcbM2/7 contain unique residues Trp32 and Thr98, whereas the corresponding residue of Trp32 is a Phe in Type I and Type II isoforms, the corresponding residue of Thr98 is a Phe in Type IV isoform, therefore exclude the possibility of all other LhcbM proteins. The density of Phe147 side-chain was used to further verify the assignment of LhcbM2/7. (D) LhcbM5 possesses the longest N-terminal region among all LhcbM proteins (underlined by a red line), which shows clear densities in the map. The assignment of LhcbM5 was further verified by the densities of specific residues Phe197 and Phe254. The former is absent, while the latter is a residue with small side-chain (Gly or Thr) in all other LhcbM proteins. (E) LhcbM2/7 contain unique residues Trp32 and Thr98, whereas the corresponding residue of Trp32 is a Phe in Type I and Type II isoforms, the corresponding residue of Thr98 is a Phe in Type IV isoform, therefore exclude the possibility of all other LhcbM proteins. The density of Phe147 side-chain was used to further verify the assignment of LhcbM2/7. (F) Type I LhcbM proteins have Phe40 and Trp48 in the N-terminal regions, whereas Type III and Type IV have Trp and Phe in the corresponding positions, therefore can be excluded. In addition, Phe117 shows clear density for its side-chain, which excludes the possibility of Type II LhcbM protein as LhcbM5 has a Ile at the corresponding position. The well-defined densities of Met213, Met216 and Thr242 further exclude LhcbM6, LhcbM9, and LhcbM8 and LhcbM9.

**Fig. S6.**
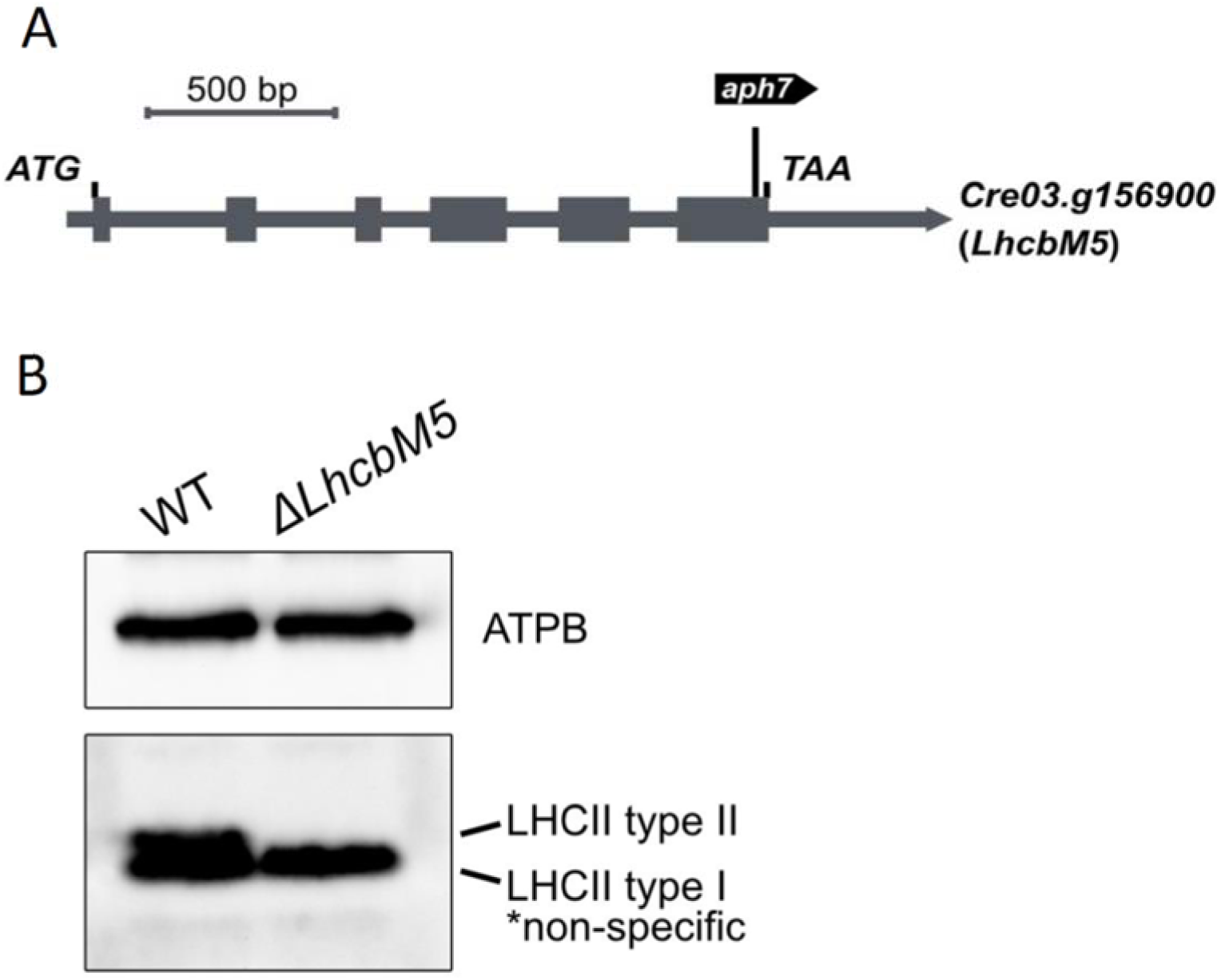
Characterization of the mutations in LhcbM5. (A) A schematic diagram of the gene structure (chromosome) of *LhcbM5* (*Cre03.g156900*) is illustrated. The translation start and stop codons and the position and orientation of the inserted a*ph7* tags in the mutant identified are all indicated. (B) Immunoblot analysis of LhcbM5. ATPB proteins were included as loading controls.

**Fig. S7.**
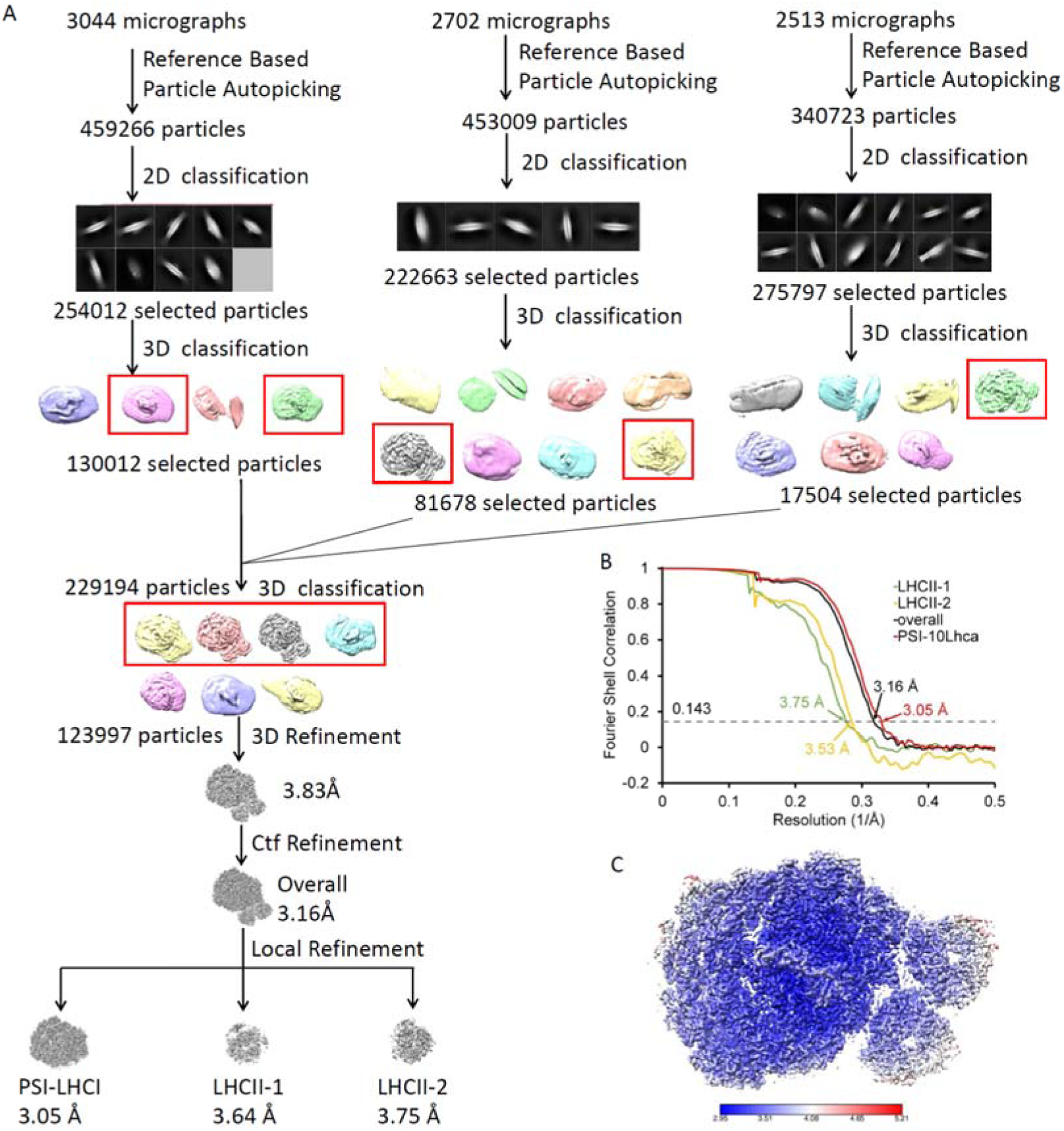
Single particle cryo-EM data processing of *Cr*PSI-LHCI-LHCII supercomplex from *ΔLhcbM1* mutant strain. (A) Single particle cryo-EM data processing procedure. (B) The gold standard Fourier shell correlation (FSC) curves of the final density map with criterion of 0.143. (C) Local resolution of the cryo-EM map estimated by ResMap.

**Fig. S8.**
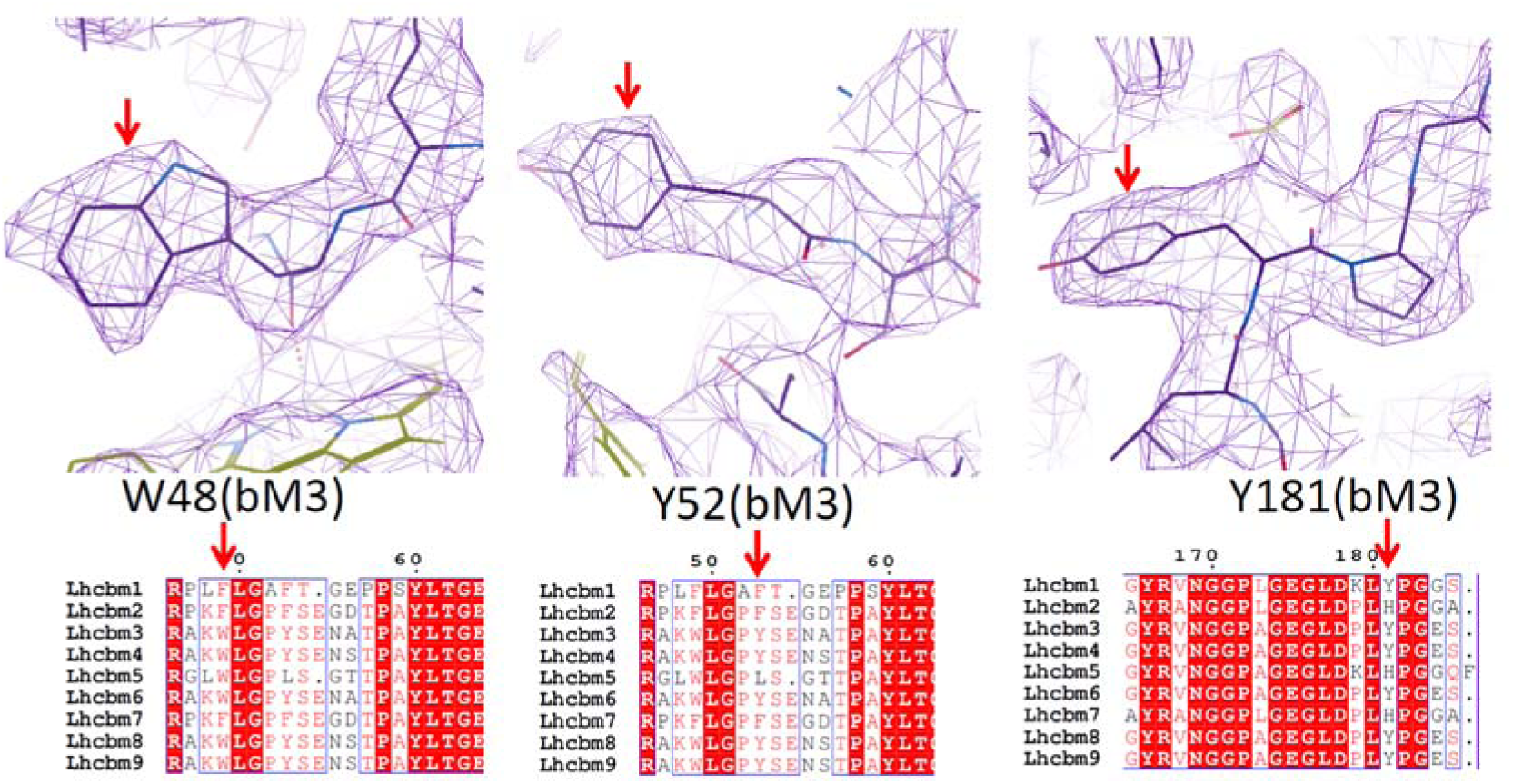
Identification of a Type I isoform replacing LhcbM1 in the structure of PSI-LHCI-LHCII supercomplex from *ΔLhcbM1* mutant strain. Map features of characteristic residues and the corresponding sequence. Type I LhcbM proteins have a Trp48 in the N-terminal regions, whereas Type III and Type IV have a Phe in the corresponding position, therefore can be excluded. Residue Tyr52 shows clear density for its side-chain, which excludes the possibility of Type II LhcbM protein as LhcbM5 has a Leu at the corresponding position. The map features of Tyr181 further verify the assignment of Type I isoform. Unique residues used for identification are indicated by red arrows.

**Fig. S9.**
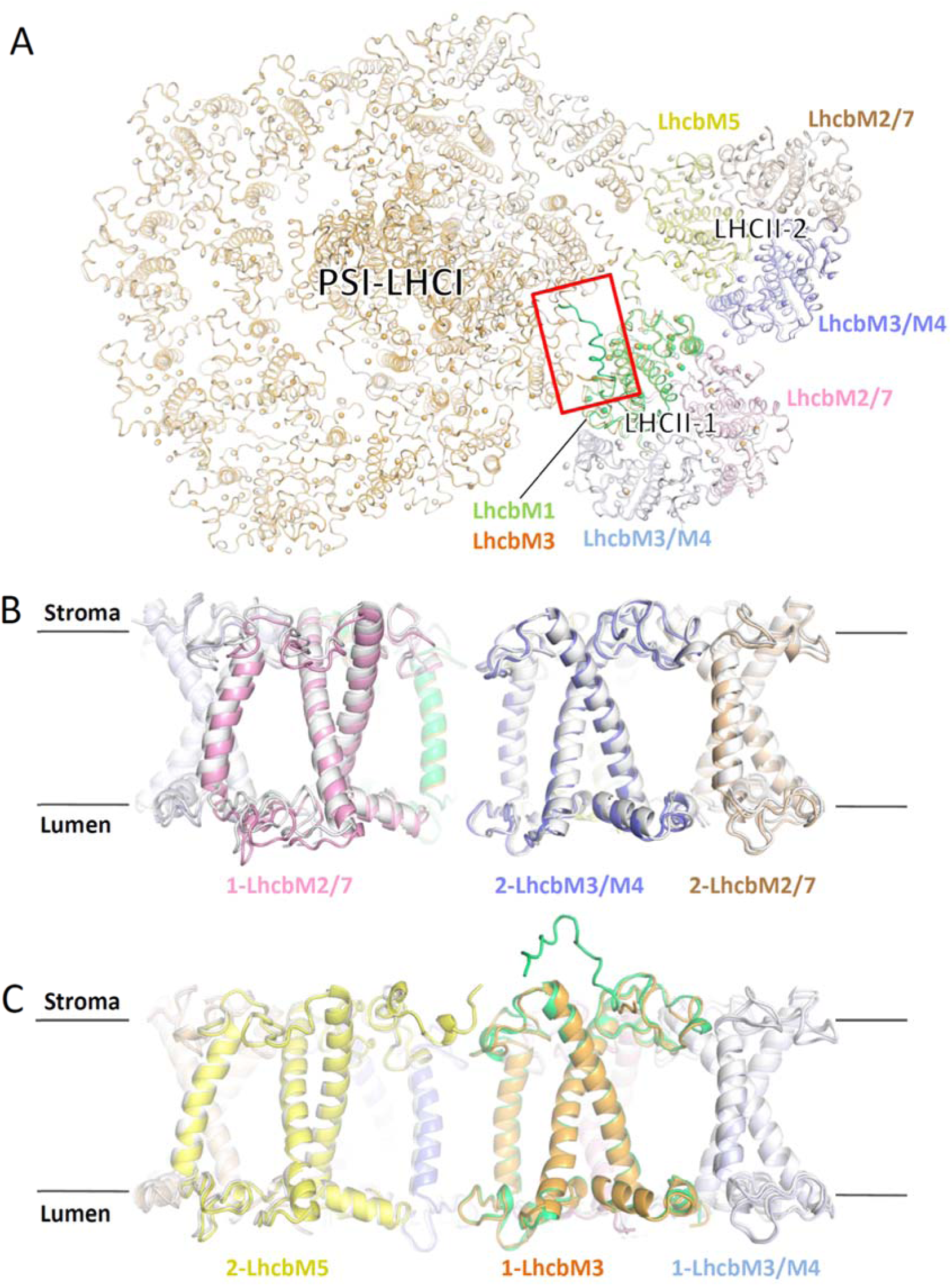
Comparison of PSI-LHCI-LHCII supercomplex from *ΔLhcbM1* mutant and in the native form. (A) Structural comparison of PSI-LHCI-LHCII supercomplex from *ΔLhcbM1* mutant and in the native form, superposed on PsaA. The native supercomplex is shown in white, with LhcbM1 of LHCII-1 highlighted in lime-green. The supercomplex from *ΔLhcbM1* mutant is colored orange for the PSI-LHCI moiety and the newly incorporated LhcbM3 of LHCII-1, while other LhcbM proteins are colored the same as in Fig. 1A. The N-terminal regions of LhcbM1 (lime-green) and the newly incorporated LhcbM3 (orange) in the two structures are highlighted by a red square. (B, C) Side view of structural comparison of two LHCII trimers in the *Cr*PSI-LHCI-LHCII supercomplex from *ΔLhcbM1* mutant and in native form, viewed from the periphery of two trimers (B) and from the PSI-LHCII interface (C). The color codes are the same as in (A). LhcbM2/7 of LHCII-1 shows the largest shift towards lumen as shown in (B).

**Fig. S10.**
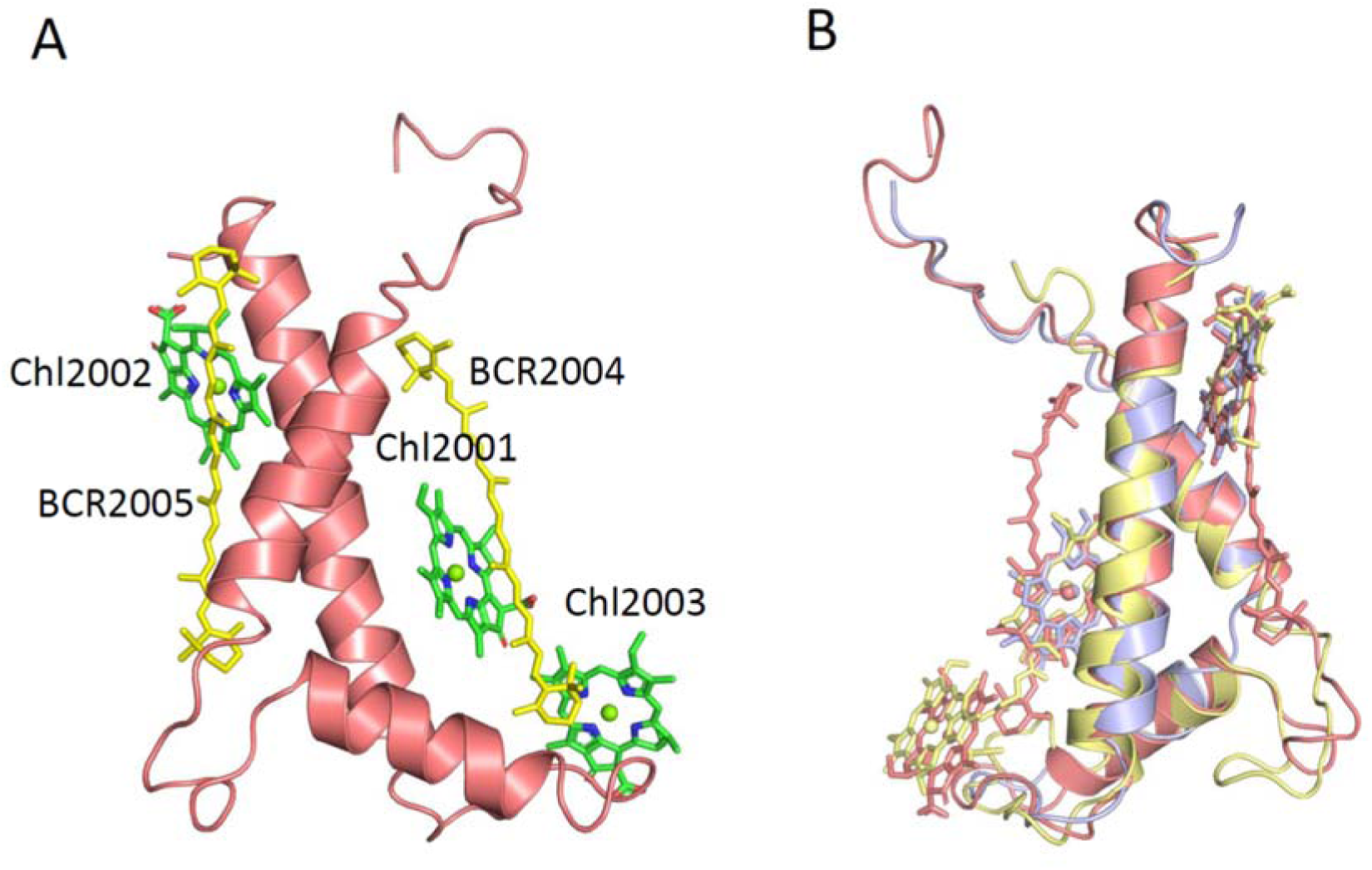
Overall structure of *Cr*PsaO. (A) Cartoon representation of *Cr*PsaO viewed from the membrane plane. *Cr*PsaO possesses two TMHs and one amphiphilic helix along the luminal surface, and binds three chlorophyll and two β-carotene molecules. The two β-carotenes are located at grooves formed by the two TMHs of PsaO from opposite sides. Three chlorophylls and two β-carotenes (BCRs) are shown in stick mode and labeled. (B) Structural comparison of *Cr*PsaO (salmon) with PsaO from maize (light-blue, PDB code 5ZJI) and red alga (yellow, PDB code 5ZGH).

**Table S1.** Cryo-EM data collection, refinement and validation statistics of *Cr*PSI-LHCI-LHCII structures.

|  | Native<br>supercomplex | Supercomplex<br>from *ΔLhcbM1*<br>mutant |
| --- | --- | --- |
| **Data collection and processing** |  |  |
| Magnification | 130,000 | 130,000 |
| Voltage (kV) | 200 | 200 |
| Electron exposure (e<sup>-</sup>/Å) | 50 | 50 |
| Defocus range ( μm ) | 1.5-2.0 | 1.5-2.2 |
| Pixel size ( Å ) | 1.00 | 1.00 |
| Symmetry imposed | C1 | C1 |
| Initial particle images (no.) | 603,892 | 1,252,998 |
| Final particle images (no.) | 55,601 | 123,997 |
| Map resolution (Å) | 2.84 | 3.16 |
| FSC threshold | 0.143 | 0.143 |
| Map resolution range (Å) | 2.65-5.58 | 2.95-5.21 |
| **Refinement** |  |  |
| Initial model used (PDB code) | 6IJO,6KAD,5ZJI | Native<br>supercomplex |
| Model resolution (Å) | 2.89 | 3.18 |
| FSC threshold | 0.5 | 0.5 |
| Map sharpening B factor (Å<sup>2</sup>) | -66.99 | -91.75 |
| Model composition |  |  |
| Nonhydrogen atoms | 69,633 | 69,450 |
| Protein residues | 5995 | 5984 |
| Ligands | 471 | 471 |
| B factor (Å<sup>2</sup>) |  |  |
| Protein | 21.36 | 44.91 |
| Ligand | 24.06 | 53.47 |
| R.m.s. deviations |  |  |
| Bond lengths (Å) | 0.009 | 0.007 |
| Bond angles (°) | 2.043 | 1.96 |
| Validation |  |  |
| MolProbity score | 2.25 | 1.93 |
| Clash score | 8.53 | 10.35 |
| Poor rotamers (%) | 3.55 | 0.19 |
| Ramachandran plot |  |  |
| Favored (%) | 94.65 | 94.07 |
| Allowed (%) | 5.24 | 5.89 |
| Disallowed (%) | 0.1 | 0.03 |

**Table S2.** Summarization of the structural model of Native *Cr*PSI-LHCI-LHCII structure.

| Subunit | Traced residues | Chlorophylls | Carotenoids | Lipids | Others |
| --- | --- | --- | --- | --- | --- |
| **PSI-LHCI** |  |  |  |  |  |
| PsaA | 741 | 45 Chl | 6 BCR | 2 PG, 1 MGDG | 1 PQN, 1 SF4, |
| PsaB | 733 | 40 Chl | 10 BCR | 1 DGDG, 1 PG | 1 PQN |
| PsaC | 80 |  |  |  | 2 SF4 |
| PsaD | 143 |  |  |  |  |
| PsaE | 63 |  |  |  |  |
| PsaF | 164 | 3 Chl | 1 BCR |  |  |
| PsaG | 94 | 2 Chl | 1 BCR |  |  |
| PsaH | 100 | 2 Chl |  | 1 PG, 1 MGDG |  |
| PsaI | 39 |  |  |  |  |
| PsaJ | 41 | 1 Chl | 1 BCR | 2 MGDG |  |
| PsaK | 86 | 4 Chl | 2 BCR |  |  |
| PsaL | 159 | 5 Chl | 4 BCR | 1 MGDG |  |
| PsaO | 97 | 3 Chl | 2 BCR | 1 PG |  |
| Lhca1<sub>in</sub> | 194 | 14 Chl | 1 Lut, 1 Vio, 1 BCR | 1 PG |  |
| Lhca1<sub>out</sub> | 194 | 14 Chl | 1 Lut, 1 Vio, 1 BCR | 1 PG |  |
| Lhca2 | 217 | 13 Chl | 1 Lut, 1 Vio, 1 BCR | 1 PG |  |
| Lhca3 | 220 | 14 Chl | 1 Lut, 1 Vio, 3 BCR | 2 PG |  |
| Lhca4 | 210 | 15 Chl | 1 Lut, 1 Vio, 1 BCR | 1 PG, 2 MGDG |  |
| Lhca5 | 227 | 17 Chl | 1 Lut, 1 Vio, 1 Neo, 1 BCR | 2 PG, 1 MGDG |  |
| Lhca6 | 230 | 17 Chl | 1 Lut, 1 Vio, 1 Neo, 1 BCR | 1 PG |  |
| Lhca7 | 213 | 15 Chl | 1 Lut, 1 Vio, 2 BCR | 1 PG |  |
| Lhca8 | 216 | 14 Chl | 1 Lut, 1 Vio, 1 BCR | 2 PG, 1 MGDG |  |
| Lhca9 | 183 | 12 Chl | 1 Lut, 1 Vio, 1 BCR | 4 PG, 1 MGDG |  |
| **LHCII-1** |  |  |  |  |  |
| LhcbM1 | 232 | 8 Chl<sub>a</sub>, 6 Chl *b* | 2 Lut, 1 Vio, 1 Neo | 1 PG |  |
| LhcbM2 | 220 | 8 Chl<sub>a</sub>, 6 Chl *b* | 2 Lut, 1 Vio, 1 Neo | 1 PG |  |
| LhcbM3 | 220 | 8 Chl<sub>a</sub>, 6 Chl *b* | 2 Lut, 1 Vio, 1 Neo | 1 PG |  |
| **LHCII-2** |  |  |  |  |  |
| LhcbM2 | 220 | 8 Chl<sub>a</sub>, 6 Chl *b* | 2 Lut, 1 Vio, 1 Neo | 1 PG |  |
| LhcbM3 | 219 | 8 Chl<sub>a</sub>, 6 Chl *b* | 2 Lut, 1 Vio, 1 Neo | 1 PG |  |
| LhcbM5 | 237 | 8 Chl<sub>a</sub>, 6 Chl *b* | 2 Lut, 1 Vio, 1 Neo | 1 PG, 1 MGDG |  |
| Total | 5992 | 334 | 87 | 39 | 12 |

BCR, β-carotene; Lut, lutein; Vio, violaxanthin; Neo, neoxanthin; PG, phosphatidyl glycerol; DGDG, digalactosyldiacyl glycerol; MGDG, monogalactosyldiacyl glycerol; PQN, phylloquinone; SF4, sulphur-iron cluster.

## Reference

1. Dekker, J.P. & Boekema, E.J. Supramolecular organization of thylakoid membrane proteins in green plants. Biochim Biophys Acta 1706, 12–39 (2005).

2. Croce, R. & van Amerongen, H. Light-harvesting and structural organization of Photosystem II: from individual complexes to thylakoid membrane. J Photochem Photobiol B 104, 142–53 (2011).

3. Pan, X., Cao, P., Su, X., Liu, Z. & Li, M. Structural analysis and comparison of light-harvesting complexes I and II. Biochim Biophys Acta Bioenerg 1861, 148038 (2020).

4. Elrad, D., Niyogi, K.K. & Grossman, A.R. A major light-harvesting polypeptide of photosystem II functions in thermal dissipation. Plant Cell 14, 1801–16 (2002).

5. Stauber, E.J. et al. Proteomics of Chlamydomonas reinhardtii light-harvesting proteins. Eukaryot Cell 2, 978–94 (2003).

6. Tokutsu, R., Kato, N., Bui, K.H., Ishikawa, T. & Minagawa, J. Revisiting the supramolecular organization of photosystem II in Chlamydomonas reinhardtii. J Biol Chem 287, 31574–81 (2012).

7. Ozawa, S.I. et al. Configuration of Ten Light-Harvesting Chlorophyll a/b Complex I Subunits in Chlamydomonas reinhardtii Photosystem I. Plant Physiology 178, 583–595 (2018).

8. Kubota-Kawai, H. et al. Ten antenna proteins are associated with the core in the supramolecular organization of the photosystem I supercomplex in Chlamydomonas reinhardtii. Journal of Biological Chemistry 294, 4304–4314 (2019).

9. Minagawa, J. & Takahashi, Y. Structure, function and assembly of Photosystem II and its light-harvesting proteins. Photosynth Res 82, 241–63 (2004).

10. Natali, A. & Croce, R. Characterization of the major light-harvesting complexes (LHCBM) of the green alga Chlamydomonas reinhardtii. PLoS One 10, e0119211 (2015).

11. Minagawa, J. State transitions—The molecular remodeling of photosynthetic supercomplexes that controls energy flow in the chloroplast. Biochimica et Biophysica Acta (BBA) - Bioenergetics 1807, 897–905 (2011).

12. Rochaix, J.D. Role of thylakoid protein kinases in photosynthetic acclimation. FEBS Lett 581, 2768–75 (2007).

13. Cariti, F. et al. Regulation of Light Harvesting in Chlamydomonas reinhardtii Two Protein Phosphatases Are Involved in State Transitions. Plant Physiol 183, 1749–1764 (2020).

14. Delosme, R., Olive, J. & Wollman, F.A. Changes in light energy distribution upon state transitions: An in vivo photoacoustic study of the wild type and photosynthesis mutants from Chlamydomonas reinhardtii. Biochimica Et Biophysica Acta-Bioenergetics 1273, 150–158 (1996).

15. Nawrocki, W.J., Santabarbara, S., Mosebach, L., Wollman, F.A. & Rappaport, F. State transitions redistribute rather than dissipate energy between the two photosystems in Chlamydomonas. Nature Plants 2(2016).

16. Nagy, G. et al. Chloroplast remodeling during state transitions in Chlamydomonas reinhardtii as revealed by noninvasive techniques in vivo. Proceedings of the National Academy of Sciences of the United States of America 111, 5042–5047 (2014).

17. Ünlü, C., Polukhina, I. & van Amerongen, H. Origin of pronounced differences in 77 K fluorescence of the green alga Chlamydomonas reinhardtii in state 1 and 2. European Biophysics Journal 45, 209–217 (2015).

18. Ünlü, C., Drop, B., Croce, R. & van Amerongen, H. State transitions in *Chlamydomonas reinhardtii* strongly modulate the functional size of photosystem II but not of photosystem I. Proceedings of the National Academy of Sciences 111, 3460–3465 (2014).

19. Pan, X. et al. Structure of the maize photosystem I supercomplex with light-harvesting complexes I and II. Science 360, 1109–1113 (2018).

20. Kargul, J. et al. Light-harvesting complex II protein CP29 binds to photosystem I of Chlamydomonas reinhardtii under State 2 conditions. FEBS Journal 272, 4797–4806 (2005).

21. Takahashi, H., Iwai, M., Takahashi, Y. & Minagawa, J. Identification of the mobile light-harvesting complex II polypeptides for state transitions in Chlamydomonas reinhardtii. Proceedings of the National Academy of Sciences 103, 477–482 (2006).

22. Drop, B., Yadav K.N, S., Boekema, E.J. & Croce, R. Consequences of state transitions on the structural and functional organization of Photosystem I in the green algaChlamydomonas reinhardtii. The Plant Journal 78, 181–191 (2014).

23. Ferrante, P., Ballottari, M., Bonente, G., Giuliano, G. & Bassi, R. LHCBM1 and LHCBM2/7 polypeptides, components of major LHCII complex, have distinct functional roles in photosynthetic antenna system of Chlamydomonas reinhardtii. J Biol Chem 287, 16276–88 (2012).

24. Lemeille, S., Turkina, M.V., Vener, A.V. & Rochaix, J.D. Stt7-dependent phosphorylation during state transitions in the green alga Chlamydomonas reinhardtii. Mol Cell Proteomics 9, 1281–95 (2010).

25. Su, X. et al. Antenna arrangement and energy transfer pathways of a green algal photosystem-I–LHCI supercomplex. Nature Plants 5, 273–281 (2019).

26. Suga, M. et al. Structure of the green algal photosystem I supercomplex with a decameric light-harvesting complex I. Nature Plants 5, 626–636 (2019).

27. Takahashi, H., Okamuro, A., Minagawa, J. & Takahashi, Y. Biochemical characterization of photosystem I-associated light-harvesting complexes I and II isolated from state 2 cells of Chlamydomonas reinhardtii. Plant Cell Physiol 55, 1437–49 (2014).

28. Pi, X. et al. Unique organization of photosystem I-light-harvesting supercomplex revealed by cryo-EM from a red alga. Proc Natl Acad Sci U S A 115, 4423–4428 (2018).

29. Liu, Z.F. et al. Crystal structure of spinach major light-harvesting complex at 2.72 angstrom resolution. Nature 428, 287–292 (2004).

30. Paulsen, H., Finkenzeller, B. & Kuhlein, N. Pigments Induce Folding of Light-Harvesting Chlorophyll Alpha/Beta-Binding Protein. European Journal of Biochemistry 215, 809–816 (1993).

31. Croce, R., Weiss, S. & Bassi, R. Carotenoid-binding sites of the major light-harvesting complex II of higher plants. Journal of Biological Chemistry 274, 29613–29623 (1999).

32. Novoderezhkin, V.I., Palacios, M.A., van Amerongen, H. & van Grondelle, R. Excitation dynamics in the LHCII complex of higher plants: modeling based on the 2.72 Angstrom crystal structure. J Phys Chem B 109, 10493–504 (2005).

33. Le Quiniou, C., van Oort, B., Drop, B., van Stokkum, I.H.M. & Croce, R. The High Efficiency of Photosystem I in the Green AlgaChlamydomonas reinhardtiiIs Maintained after the Antenna Size Is Substantially Increased by the Association of Light-harvesting Complexes II. Journal of Biological Chemistry 290, 30587–30595 (2015).

34. Lunde, C., Jensen, P.E., Haldrup, A., Knoetzel, J. & Scheller, H.V. The PSI-H subunit of photosystem I is essential for state transitions in plant photosynthesis. Nature 408, 613–615 (2000).

35. Jensen, P.E., Haldrup, A., Zhang, S. & Scheller, H.V. The PSI-O subunit of plant photosystem I is involved in balancing the excitation pressure between the two photosystems. J Biol Chem 279, 24212–7 (2004).

36. Zhang, S.P. & Scheller, H.V. Light-harvesting complex II binds to several small Subunits of photosystem I. Journal of Biological Chemistry 279, 3180–3187 (2004).

37. Suga, M. & Shen, J.R. Structural variations of photosystem I-antenna supercomplex in response to adaptations to different light environments. Current Opinion in Structural Biology 63, 10–17 (2020).

38. Kim, E., Kawakami, K., Sato, R., Ishii, A. & Minagawa, J. Photoprotective Capabilities of Light-Harvesting Complex II Trimers in the Green Alga Chlamydomonas reinhardtii. Journal of Physical Chemistry Letters 11, 7755–7761 (2020).

39. Shen, L. et al. Structure of a C2S2M2N2-type PSII-LHCII supercomplex from the green alga Chlamydomonas reinhardtii. Proc Natl Acad Sci U S A 116, 21246–21255 (2019).

40. Sheng, X. et al. Structural insight into light harvesting for photosystem II in green algae. Nat Plants 5, 1320–1330 (2019).

41. Tokutsu, R., Fujimura-Kamada, K., Yamasaki, T., Matsuo, T. & Minagawa, J. Isolation of photoprotective signal transduction mutants by systematic bioluminescence screening in Chlamydomonas reinhardtii. Sci Rep 9, 2820 (2019).

42. Depege, N., Bellafiore, S. & Rochaix, J.D. Role of chloroplast protein kinase Stt7 in LHCII phosphorylation and state transition in Chlamydomonas. Science 299, 1572–1575 (2003).

43. Lemeille, S. et al. Analysis of the Chloroplast Protein Kinase Stt7 during State Transitions. Plos Biology 7, 664–675 (2009).

44. Takahashi, H., Clowez, S., Wollman, F.A., Vallon, O. & Rappaport, F. Cyclic electron flow is redox-controlled but independent of state transition. Nat Commun 4, 1954 (2013).

45. Allorent, G. et al. A Dual Strategy to Cope with High Light in Chlamydomonas reinhardtii. The Plant Cell 25, 545–557 (2013).

46. Gorman, D.S. & Levine, R.P. Cytochrome f and plastocyanin: their sequence in the photosynthetic electron transport chain of Chlamydomonas reinhardi. Proc Natl Acad Sci U S A 54, 1665–9 (1965).

47. Iwai, M. et al. Isolation of the elusive supercomplex that drives cyclic electron flow in photosynthesis. Nature 464, 1210–3 (2010).

48. Watanabe, A., Kim, E., Burton-Smith, R.N., Tokutsu, R. & Minagawa, J. Amphipol-assisted purification method for the highly active and stable photosystem II supercomplex of Chlamydomonas reinhardtii. FEBS Letters 593, 1072–1079 (2019).

49. Chua, N.H. & Bennoun, P. Thylakoid membrane polypeptides of Chlamydomonas reinhardtii: wild-type and mutant strains deficient in photosystem II reaction center. Proc Natl Acad Sci U S A 72, 2175–9 (1975).

50. Wei, X. et al. Structure of spinach photosystem II-LHCII supercomplex at 3.2 A resolution. Nature 534, 69–74 (2016).

51. Farber, A., Young, A.J., Ruban, A.V., Horton, P. & Jahns, P. Dynamics of Xanthophyll-Cycle Activity in Different Antenna Subcomplexes in the Photosynthetic Membranes of Higher Plants (The Relationship between Zeaxanthin Conversion and Nonphotochemical Fluorescence Quenching). Plant Physiol 115, 1609–1618 (1997).

52. Mastrobuoni, G. et al. Proteome dynamics and early salt stress response of the photosynthetic organism Chlamydomonas reinhardtii. BMC Genomics 13, 215 (2012).

53. Tokutsu, R., Iwai, M. & Minagawa, J. CP29, a Monomeric Light-harvesting Complex II Protein, Is Essential for State Transitions in Chlamydomonas reinhardtii. Journal of Biological Chemistry 284, 7777–7782 (2009).

54. Mastronarde, D.N. Automated electron microscope tomography using robust prediction of specimen movements. Journal of Structural Biology 152, 36–51 (2005).

55. Wu, C., Huang, X., Cheng, J., Zhu, D. & Zhang, X. High-quality, high-throughput cryo-electron microscopy data collection via beam tilt and astigmatism-free beam-image shift. J Struct Biol 208, 107396 (2019).

56. Zheng, S.Q. et al. MotionCor2: anisotropic correction of beam-induced motion for improved cryo-electron microscopy. Nat Methods 14, 331–332 (2017).

57. Rohou, A. & Grigorieff, N. CTFFIND4: Fast and accurate defocus estimation from electron micrographs. J Struct Biol 192, 216–21 (2015).

58. Zivanov, J. et al. New tools for automated high-resolution cryo-EM structure determination in RELION-3. Elife 7(2018).

59. Pettersen, E.F. et al. UCSF Chimera--a visualization system for exploratory research and analysis. J Comput Chem 25, 1605–12 (2004).

60. Kucukelbir, A., Sigworth, F.J. & Tagare, H.D. Quantifying the local resolution of cryo-EM density maps. Nat Methods 11, 63–5 (2014).

61. Emsley, P., Lohkamp, B., Scott, W.G. & Cowtan, K. Features and development of Coot. Acta Crystallogr D Biol Crystallogr 66, 486–501 (2010).

62. Adams, P.D. et al. PHENIX: a comprehensive Python-based system for macromolecular structure solution. Acta Crystallogr D Biol Crystallogr 66, 213–21 (2010).

63. Chen, V.B. et al. MolProbity: all-atom structure validation for macromolecular crystallography. Acta Crystallogr D Biol Crystallogr 66, 12–21 (2010).

64. Elrad, D. & Grossman, A.R. A genome’s-eye view of the light-harvesting polypeptides of Chlamydomonas reinhardtii. Curr Genet 45, 61–75 (2004).

